# Validating the Cyc2 neutrophilic Fe oxidation pathway using meta-omics of Zetaproteobacteria iron mats at marine hydrothermal vents

**DOI:** 10.1101/722066

**Authors:** Sean M. McAllister, Shawn W. Polson, David A. Butterfield, Brian T. Glazer, Jason B. Sylvan, Clara S. Chan

## Abstract

Zetaproteobacteria create extensive iron (Fe) oxide mats at marine hydrothermal vents, making them an ideal model for microbial Fe oxidation at circumneutral pH. Comparison of neutrophilic Fe-oxidizer isolate genomes has revealed a hypothetical Fe oxidation pathway, featuring a homolog of the Fe oxidase Cyc2 from *Acidithiobacillus ferrooxidans*. However, Cyc2 function is not well verified in neutrophilic Fe-oxidizers, particularly in Fe-oxidizing environments. Toward this, we analyzed genomes and metatranscriptomes of Zetaproteobacteria, using 53 new high-quality metagenome assembled genomes reconstructed from Fe mats at Mid-Atlantic Ridge, Mariana Backarc, and Loihi Seamount (Hawaii) hydrothermal vents. Phylogenetic analysis demonstrated conservation of Cyc2 sequences among most neutrophilic Fe-oxidizers, suggesting a common function. We confirmed the widespread distribution of *cyc2* and other model Fe oxidation pathway genes across all represented Zetaproteobacteria lineages. High expression of these genes was observed in diverse Zetaproteobacteria under multiple environmental conditions, and in incubations. The putative Fe oxidase gene, *cyc2*, was highly expressed *in situ*, often as the top expressed gene. The *cyc2* gene showed increased expression in Fe(II)-amended incubations, with corresponding increases in carbon fixation and central metabolism gene expression. These results substantiate the Cyc2-based Fe oxidation pathway in neutrophiles and demonstrate its significance in marine Fe-mineralizing environments.

## Introduction

Neutrophilic Fe-oxidizing microbes are common in marine and terrestrial environments [1], precipitating reactive Fe oxyhydroxides that sequester organic carbon, phosphate, arsenic, and many other metals [2–4]. However, it has been difficult to study the effects of neutrophilic Fe oxidation in natural systems due to myriad challenges that have slowed the discovery of genetic markers of neutrophilic Fe oxidation. These Fe-oxidizers are difficult to culture, so only recently have we obtained enough isolate genomes to deduce hypothetical neutrophilic Fe-oxidizing pathways. Comparative genomics has led to multiple proposed pathways, each involving an outer membrane cytochrome [5–7]. However, only one pathway is present in all well-established neutrophilic Fe-oxidizing isolates (Zetaproteobacteria and Gallionellaceae), centering on a fused cytochrome-porin Cyc2 [7–10]. Yet, beyond comparative genomics, we lack evidence of the Cyc2 pathway function in neutrophilic Fe-oxidizers, particularly the uncultured Fe-oxidizers that dominate natural Fe systems.

The Cyc2 pathway in the neutrophilic Fe-oxidizers is modeled after the Fe oxidation pathways found in acidophilic Fe-oxidizers *Acidithiobacillus ferrooxidans* and *Leptospirillum* sp., where Cyc2 Fe oxidase function has been verified [11, 12]. Weak homologs to *cyc2* from these organisms were first found in the genomes of the Gallionellaceae *Sideroxydans lithotrophicus* ES-1 and *Gallionella capsiferriformans* ES-2 [13]. The genome of Zetaproteobacteria type strain *Mariprofundus ferrooxydans* PV-1, on the other hand, lacked homologs to known Fe oxidation genes until a proteome study discovered that *cyc2* was in fact encoded by PV-1 but missing from the draft genome [14]. Subsequently, *cyc2* homologs were found within the few Zetaproteobacteria lineages with genomic representation [9, 15]. Despite the identification of *cyc2*-like genes, low amino acid sequence homology (20% sequence identity between PV-1 and *A. ferrooxidans* Cyc2) suggests that their function is speculative and needs to be validated. Without a means of testing this function biochemically or genetically, we focus on more comprehensive comparative genomics and expression in Fe-oxidizing environments.

To this end, we turned to Zetaproteobacteria in natural Fe microbial mats. The Zetaproteobacteria discovered to date are all considered to be Fe-oxidizers, since every isolate grows by Fe oxidation and uncultured Zetaproteobacteria are typically found in Fe-oxidizing environments [10, 16–21]. The Zetaproteobacteria are often the dominant organisms in marine hydrothermal Fe mats [22–25], where they play a key role in mat formation [26]. This abundance and ubiquity in Fe-oxidizing mats makes Zetaproteobacteria ideal for study through metagenomic and metatranscriptomic approaches. Furthermore, their taxonomic diversity allows for a robust comparative genomics study. We sampled paired metagenomes and metatranscriptomes from three hydrothermal venting regions: Loihi Seamount, the Mid-Atlantic Ridge (Rainbow, TAG, and Snake Pit vents), and the Mariana Backarc (Urashima vent field). Recovery of high quality metagenome assembled genomes (MAGs) allowed us to improve the limited genomic representation of the Zetaproteobacteria (see [10] for a summary). Using the MAGs for reference mapping, we explored *in situ* environmental expression of the Zetaproteobacteria within undisturbed natural Fe mats. In addition, we examined expression responses to Fe(II) using shipboard Fe(II) amendment experiments. With these results, we assess and update the model neutrophilic Fe oxidation pathway expressed in natural environments.

## Materials and Methods

### Biological sample collection

Samples were collected on three separate cruises to the Mid-Atlantic Ridge (2012), Loihi Seamount (2013), and the Southern Mariana Trough (2014) (**Supplemental Table 1**). To preserve *in situ* expression, 18 samples were collected using devices half-filled with 2x RNALater (Invitrogen, Carlsbad, CA, USA). Samples collected using a syringe sampler [27] provided ~10-30 mL of mat material representing a discrete microbial population, as opposed to 2 L (scoop) or > 5 L (suction sample) of bulk sample. After settling for a few hours at 4°C, samples were stored at −80°C.

### Geochemical measurements and sampling

At the Mid-Atlantic Ridge and Loihi Seamount, geochemistry was measured *in situ* using cyclic voltammetry, as described in MacDonald et al. [28] (MAR) and Chan et al. [26] (Loihi). The detection limits were 3 µM O_2_, 7–10 µM Fe^2+^, and 0.1 µM sulfide [28, 29].

At Mariana, geochemistry samples were collected using the Hydrothermal Fluid and Particle Sampler (HFPS) [30] or the Microbial Mat sampler [27]. The HFPS pulls fluid through a titanium inlet nozzle at 1-4 L/min. The fluid flows through a continuously flushed titanium and Teflon manifold and is diverted into sample containers or to a SeaBird (Bellevue, WA) SBE 63 oxygen sensor and an AMT (Rostock, Germany) deep-sea glass pH electrode. In extremely low outflow vent environments, seawater will be entrained in the HFPS and dilute the *in situ* fluid. We collected temperature, pH, and O_2_ concentrations for ambient water, at the surface of the chimney, and with the nozzle inserted into the microbial mat. HFPS chemistry results represent the fluid composition at the measured temperature, including any entrained seawater that occurs during sampling. The Microbial Mat sampler has a much lower intake rate (< 0.2 L min^−1^) and is better able to capture chemical micro-environments. The microbial mat sampler was equipped with 0.22 µm in-line filtering and a check valve. Fe(II) and total Fe concentrations were assayed using the ferrozine method with 40 mM sulfamic acid to stabilize Fe(II) [31, 32]; detection limit estimated at 0.12 µM Fe(II). Samples recovered with the HFPS were processed as described previously [30], analyzed shipboard for pH by glass electrode, and on shore for total dissolved iron by atomic absorption at NOAA/PMEL and by ICPMS at the University of Washington Department of Civil Engineering.

### Fe(II) amendment experiments

Shipboard Fe(II) amendment experiments were conducted on bulk mat samples from Loihi (J2-677-SSyellow) and Mariana (J2-801-SC8). Samples were transported to the ship after 2 hrs (Loihi) and 11 hrs (Mariana) of ROV operations, and allowed to settle at 4°C for 1 hr prior to starting the experiment. One sample was taken immediately prior to Fe(II) amendment (Pre-Fe(II) addition). Water bath temperature and initial Fe(II) amendment concentration were set to mimic environmental conditions. Fe oxidation pseudo-first-order rate constants (*k_1_*) were calculated using a log-linear fitted trend line.

At Loihi Seamount, Fe mat floc was added to two 250 mL vessels, one remained alive while the other was killed using 1 mM azide. Both vessels were shaken by hand in a 35°C water bath. To initiate the experiment, 100 µM FeCl_2_ was added. After this, starting after 2 min and subsequently at 10 min intervals, samples from each vessel were removed for Fe(II) and total Fe measurements by the ferrozine method [31]. Concurrently, 30 mL from the living vessel was mixed 1:1 with 2x RNALater (Invitrogen). This mixture was held at 4°C for a few hours prior to freezing at −80°C.

At Mariana, Fe mat material was sparged with a 2% O_2_ gas mix (pH 5.9/6.2 before/after sparge). Each time point (*n* = 5) and treatment (duplicate living and 3 mM azide-killed) had its own 125 mL reaction vessel with 30 mL mat material. In addition to a Pre-Fe(II) addition sample, one sample was taken at the end which did not experience any Fe(II) addition. Both of these non-amended samples had low Fe(II) concentrations (BD and 0.3 µM, respectively). Each reaction vessel was amended with 333 µM FeCl_2_, and suspended in a 28°C water bath with mixing by hand. Starting after 4-6 min and subsequently at 10 min intervals, one vessel was sacrificed at each time point, sampling for Fe(II), total Fe, pH, and mixing 25 mL of material 1:1 with 2x RNALater.

### DNA and RNA extraction

DNA samples were extracted using the FastDNA SPIN kit for soil (MP Biomedicals, Santa Ana, CA, USA) following the manufacturer’s instructions, except that 250 µL of a 0.5 mM sodium citrate solution (pH 5.8) was added prior to lysis. RNA samples were extracted using the NucleoSpin RNA kit (Macherey-Nagel, Bethlehem, PA, USA), with modifications detailed in supplemental methods. RNA extractions were used for metatranscriptome library prep after non-degraded total RNA (visible 16S and 23S rRNA peaks) was confirmed by Fragment Analyzer (2.2-9.3 RQN; median 5.7 RQN) (Advanced Analytical, Ankeny, IA, USA).

### 16S rRNA, metagenome, and metatranscriptome sequencing

Microbial community composition was first estimated using a PacBio-based 16S rRNA gene survey (see supplemental methods). Zetaproteobacteria operational taxonomic units (ZOTUs) were classified from these 16S rRNA gene sequences using ZetaHunter [33]. Samples were chosen for metagenomic (MG) and metatranscriptomic (MT) sequencing based on the microbial community composition and Zetaproteobacteria diversity. MG and MT libraries were prepped and sequenced at the University of Delaware Sequencing and Genotyping Center. Sequencing details are provided in the supplemental methods.

### Metagenome assembly, binning, and annotation

Raw sequence reads were trimmed to remove adaptors, poor quality regions, and short sequences (Trimmomatic) [34] and paired reads were merged if overlapping (Flash) [35]. Metagenome libraries were assembled from QC’ed reads using metaSPAdes v3.10, with read error correction disabled to improve recovery of real community genomic variation [36]. Only contigs mapping ≥1x read coverage over 90% of their length were utilized in downstream analysis (~92% remained).

Metagenome assembled genomes (MAGs) were binned using four binning programs: MaxBin [37], MetaBAT (super specific & very sensitive settings) [38], CONCOCT [39], and BinSanity [40]. The resulting bins were combined and dereplicated using DAS Tool [41]. Manual taxonomic and outlier (GC/coverage) curation of bins was performed in ggkbase (ggkbase.berkeley.edu), with additional curation performed using Anvi’o v3 [42]. Finalized curated bins were tested for completeness and redundancy using CheckM [43], classified using PhyloSift [44], and gene calling and SEED annotation was performed using RAST [45]. RAST gene calls were used for COG annotation within Anvi’o [42, 46] and KEGG annotation through BlastKoala [47]. Genes of interest (e.g. *cyc2*, *cyc1*, and terminal oxidases) were further manually curated based on evidence using NCBI BLASTp [48] against Zetaproteobacteria protein references. Gene annotation was assessed with maximum likelihood phylogenetic trees built from alignments using RAxML [49]. The Cyc2 phylogenetic tree was constructed from an alignment of 634 unique Cyc2 protein sequences identified from NCBI and IMG databases using BLASTp [48, 50, 51]. Additional information on the Cyc2 sequences and tree construction are provided in the supplemental methods.

### RNA read recruitment and expression estimates

Raw total RNA reads were quality controlled (see above) using Trimmomatic (average 99% of reads passed) [34]. Ribosomal RNA reads were removed using SortMeRNA (v2.1b) [52]. The resulting non-rRNA reads, primarily mRNA, were used for subsequent recruitment for expression estimates. MT reads were recruited to the MG from the same sample, with the exceptions: 665-MMA4 was recruited to 665-MMA12; S7_B5, S8_B2, S8_B3, S9 and S24 were recruited to S7_B4 MG. Reads were mapped using bowtie2, with default parameters [53].

To determine gene read recruitment, we used BEDTools to extract the read count from each gene coordinate region [54]. We used three normalization methods for estimating gene expression: 1) transcripts per million (TPM), normalizing for sequencing effort and gene and read lengths [55]; 2) TPM values further normalized to the average expression of six constitutively expressed genes (*adk*, *gyrA*, *recA*, *rpoB*, *rpoC*, *secA*) [56] to correct for changes in organism relative abundance (constitutive normalized expression); 3) TPM values normalized to the maximum expression for the time series for visual representation.

### Data accessibility

High quality full-length reads (20-pass minimum) from the PacBio 16S rRNA gene survey were submitted to GenBank (MK048478-MK048944). Raw metagenome and metatranscriptome reads, as well as 5-pass-filtered PacBio 16S rRNA gene reads, were submitted to the NCBI SRA under BioProject accession PRJNA555820. Metagenome assemblies from this study and reassembled metagenome assemblies from Fullerton et al. [9] were submitted to the JGI IMG database (Sequence Project IDs Gp0295814-Gp0295821, Gp0295823 [this study]; Analysis Project IDs Ga0256915, Ga0257019-Ga0257023 [Fullerton et al. [9]]). Zetaproteobacteria MAGs were also submitted to the JGI IMG database (see sequence project IDs listed above). Specific accession numbers per sample are shown in **Supplemental Table 2**.

## Results

### Microbial Fe mat sampling and geochemistry

Over three expeditions, we sampled a wide diversity of Fe microbial mats (**Supplemental Table 1**). Sampled mats varied in their physical setting, with meter-scale Loihi mats found in direct and indirect flow from vent fissures, tens of cm-scale mat mounds at the MAR at the diffuse-venting periphery of black smoker fields, and the mats at the Mariana Backarc Urashima vent fields covering a seven meter-tall Fe chimney (**Supplemental Figure 1; Supplemental text)**. Temperatures ranged from 10-63°C, while geochemical conditions also varied widely, notably concentrations of Fe(II) from 1.3-190 µM and O_2_ from <3-123 µM within the mats (Table 1). Mariana mats had shallow O_2_ gradients while at Loihi, O_2_ was undetectable (<3 µM) at 1 cm below the mat surface. These Fe(II) and O_2_ conditions favor biotic Fe oxidation [10]. At Mariana, total dissolved Fe was depleted by 49%-74% in our low temperature mats relative to the conservative mixing of the local high-temperature zero-Mg endmember (**Supplemental Figure 2**), which suggests that a substantial amount of Fe is being oxidized and precipitated within these mats.

**Table 1.**
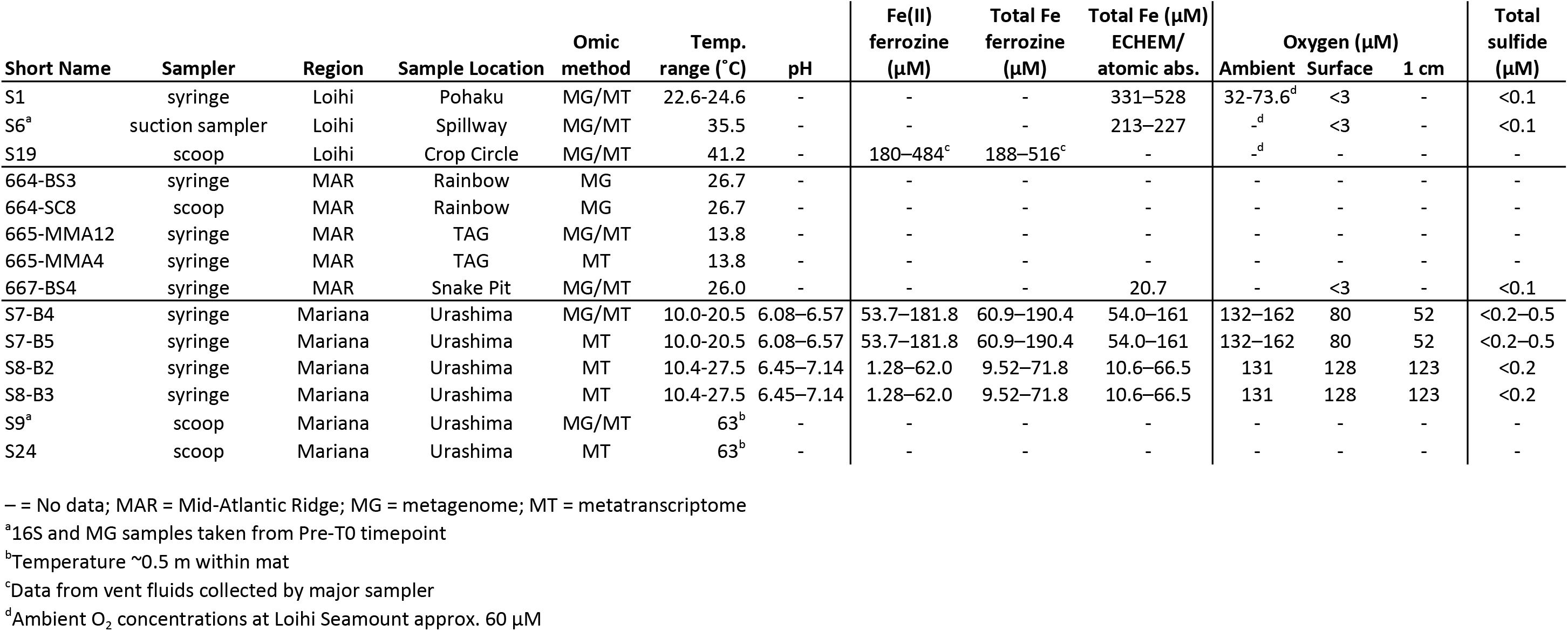
Summary of key geochemistryfor each sample.

### Zetaproteobacteria abundance and diversity

We initially assessed Zetaproteobacteria abundance and diversity using a 16S rRNA gene survey (**Supplemental Figure 3**). Fe mat communities at all sites hosted abundant Zetaproteobacteria populations, from 16.4 to 95.9% of the total bacterial community at Loihi, 10.7 to 31.3% at the Mid-Atlantic Ridge, and 37.1 to 79.9% at Mariana. Many samples were dominated by the Zetaproteobacteria, notably sample S1 (96% Zetaproteobacteria), a cm-scale sample of actively growing Fe mat surface. In addition to Zetaproteobacteria, the mats hosted variable flanking microbial communities that differed between the three sites (**Supplemental Figure 3**; **Supplemental Figure 4**), but were similar to previous studies [9, 22, 23, 57, 58]. Overall, the relatively simple, Zetaproteobacteria-rich composition of these marine Fe mats makes them good systems for studying neutrophilic Fe oxidation mechanisms.

We used the 16S rRNA gene community profiling results to choose MG and MT samples, aiming to recover high abundance and diverse Zetaproteobacteria to produce high quality genomes with sufficient MT read depth (**Supplemental Figure 3**). We recovered 126 total high quality MAGs from our samples (>70% complete, <10% redundant) (**Supplemental Table 2; Supplemental Table 3**) along with 79 improved MAGs by reanalyzing a Loihi metagenomic dataset by Fullerton et al. [9] (**Supplemental Table 3; Supplemental methods**). Of these, 53 MAGs belonged to the Zetaproteobacteria (selected genomes in **Supplemental Table 4**), which were compared to a collection of published high quality genomes (**Supplemental methods**).

MAGs from this study improve the representation of nine different ZOTUs spanning the Zetaproteobacteria phylogenetic tree by providing 2 to 13 additional high quality MAGs for each of these ZOTUs (Figure 1; **Supplemental Table 4**). Many of these ZOTUs previously had poor genome representation (see labeled ZOTUs in Figure 1B). These diverse ZOTUs were abundant and active within our Fe mats (abundance by 16S rRNA gene and MG; activity by MT) (Figure 2). MAG relative abundance generally matched relative activity, with the exception of MAG S6_Zeta1 (ZOTU6), which had higher activity than expected, likely in response to the shipboard incubation conditions. By substantially improving Zetaproteobacteria genome representation and pairing this with metatranscriptomes, we are poised to investigate genetic commonalities and diversity across the Zetaproteobacteria, particularly of the Fe oxidation mechanism.

**Figure 1.**
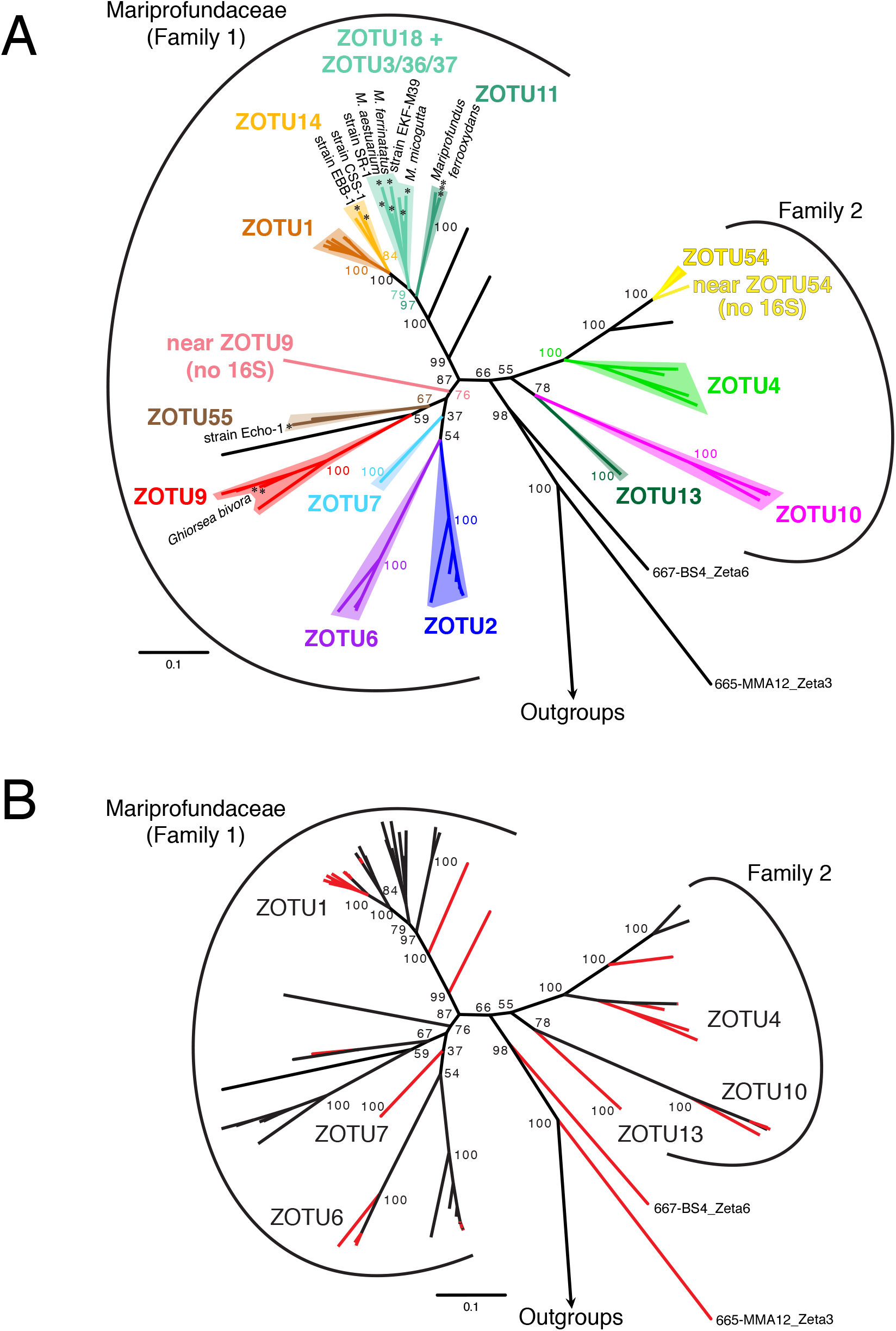
Zetaproteobacteria concatenated ribosomal protein reference maximum likelihood tree (100 bootstraps) showing (A) the most commonly sampled ZOTUs and (B) highlighting genomes produced by this study. (A) All isolates of the Zetaproteobacteria are marked with an asterisk. Deep branching genomes 665-MMA12_Zeta3 and 667-BS4_Zeta6 were classified as Zetaproteobacteria, though are deeper rooted than any prior lineage. (B) Genomes produced by this study are highlighted in red. Six ZOTUs that lacked sufficient depth for comparative genomics prior to this study are labeled.

**Figure 2.**
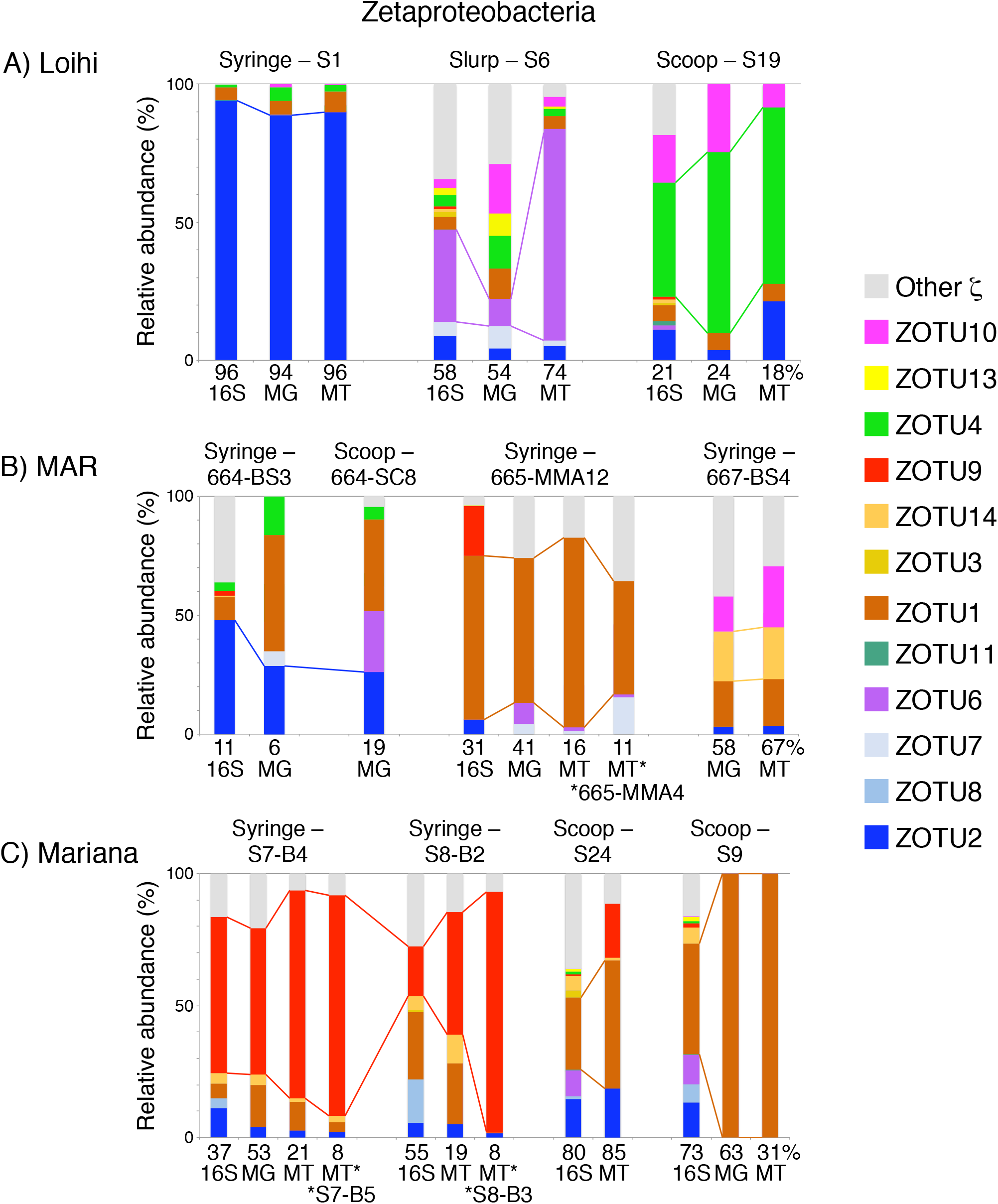
Comparison of 16S rRNA gene, metagenome (MG), and metatranscriptome (MT) relative abundance for the Zetaproteobacteria at Loihi Seamount (A), the Mid-Atlantic Ridge (B), and Southern Mariana Trough (C). The relative abundance of the most abundant Zetaproteobacteria operational taxonomic unit (ZOTU) by 16S rRNA gene are tracked across similar Fe mat samples from the same region. Asterisks denote MT from different samples that were mapped to the indicated MG. Percentages are shown at the bottom of each bar graph to indicate the relative proportion of Zetaproteobacteria in each sample (see Supplemental Figure 4).

### Phylogeny of the putative Fe oxidase Cyc2

The key component of the proposed neutrophilic Fe oxidation pathway is Cyc2, which has been shown to oxidize Fe(II) in acidophiles. Our preliminary analyses showed that some Zetaproteobacteria genomes have multiple *cyc2* copies that were not closely related. To investigate these, we developed a comprehensive Cyc2 phylogeny. This phylogeny includes sequences from terrestrial to marine, circumneutral to acidic environments, as well as both known Fe-oxidizers and organisms not known to oxidize Fe (Figure 3; see **Supplemental Figure 5** for detailed tree with sequence names). Cyc2 sequences form three clusters, but the Fe-oxidizing function has only been verified for the Cluster 2 *Acidithiobacillus ferrooxidans* Cyc2 [11] and the Cluster 3 Cyc2 homolog Cyt_572_ of *Leptospirillum* sp. [12]. However, most of the neutrophilic Fe-oxidizers fall within Cluster 1, a well-supported group (93% bootstrap) that is largely comprised of the Zetaproteobacteria, Gallionellaceae, and *Chlorobium ferrooxidans*. This strongly suggests that Cluster 1 Cyc2 share a common function.

**Figure 3.**
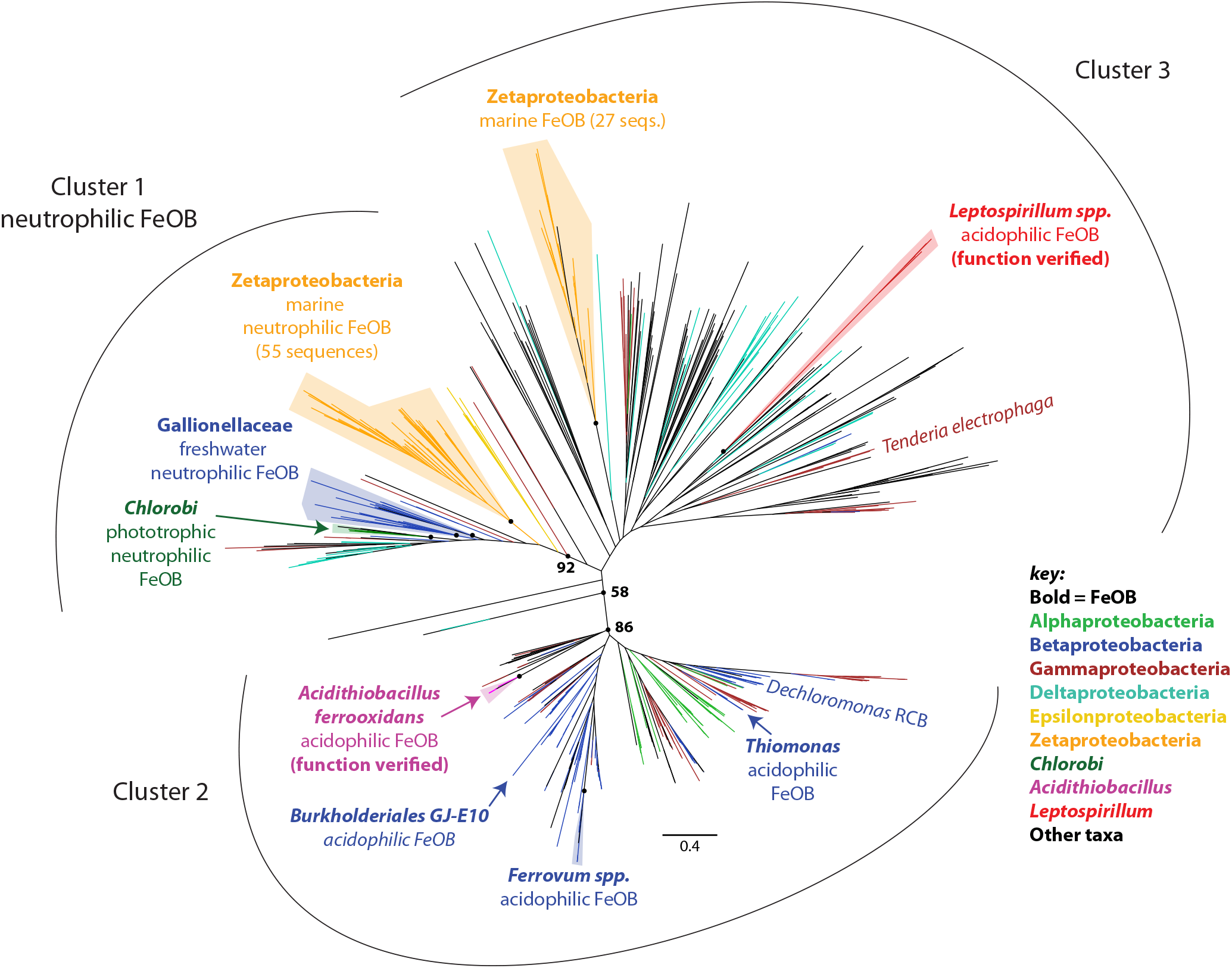
Cyc2 maximum likelihood phylogenetic tree (100 bootstraps), showing three distinct clusters. All groups of Fe-oxidizing bacteria are labeled, in addition to the electrode-oxidizing *Tenderia electrophaga*. Zetaproteobacteria and Gallionellaceae cluster with other neutrophilic Fe oxidizers in Cluster 1 (92% cluster support). Fe oxidation has been demonstrated for Cyc2 from Cluster 2 *Acidithiobacillus ferrooxidans* and Cluster 3 *Leptospirillum ferriphilum*. Unlabeled circles at nodes correspond to the following bootstrap values: Zetaproteobacteria – cluster 1 (97%), Gallionellaceae (87%/63%), Chlorobi (99%), *Acidithiobacillus ferrooxidans* (100%), *Ferrovum* spp. (100%), Zetaproteobacteria – cluster 3 (99%), *Leptospirillum* spp. (100%).

### Putative Fe oxidation pathway distribution revealed by comparative genomics of the Zetaproteobacteria

Our Cyc2 phylogeny shows that Zetaproteobacteria possess Cyc2 from both Clusters 1 and 3. All ZOTUs possess a Cluster 1 *cyc2* gene; with our new genomes, this includes four additional ZOTUs that are now known to possess *cyc2* (ZOTUs 1, 7, 13, and 14; Figure 4). In contrast, fewer ZOTUs have the Cluster 3 *cyc2* gene, and 65% of genomes with Cluster 3 *cyc2* (*n* = 15) also have Cluster 1 *cyc2*. This suggests both Cyc2 types have a use in the Zetaproteobacteria, though it is unknown how Cluster 1 and 3 Cyc2 may differ in function. ZOTU2 is unusual in that only 3 of the 10 genomes appear to have *cyc2*, though this may be due to assembly issues specific to ZOTU2 (**Supplemental text**). In any case, the presence of *cyc2* in all Zetaproteobacteria OTUs suggests its centrality to these neutrophilic Fe-oxidizers.

**Figure 4.**
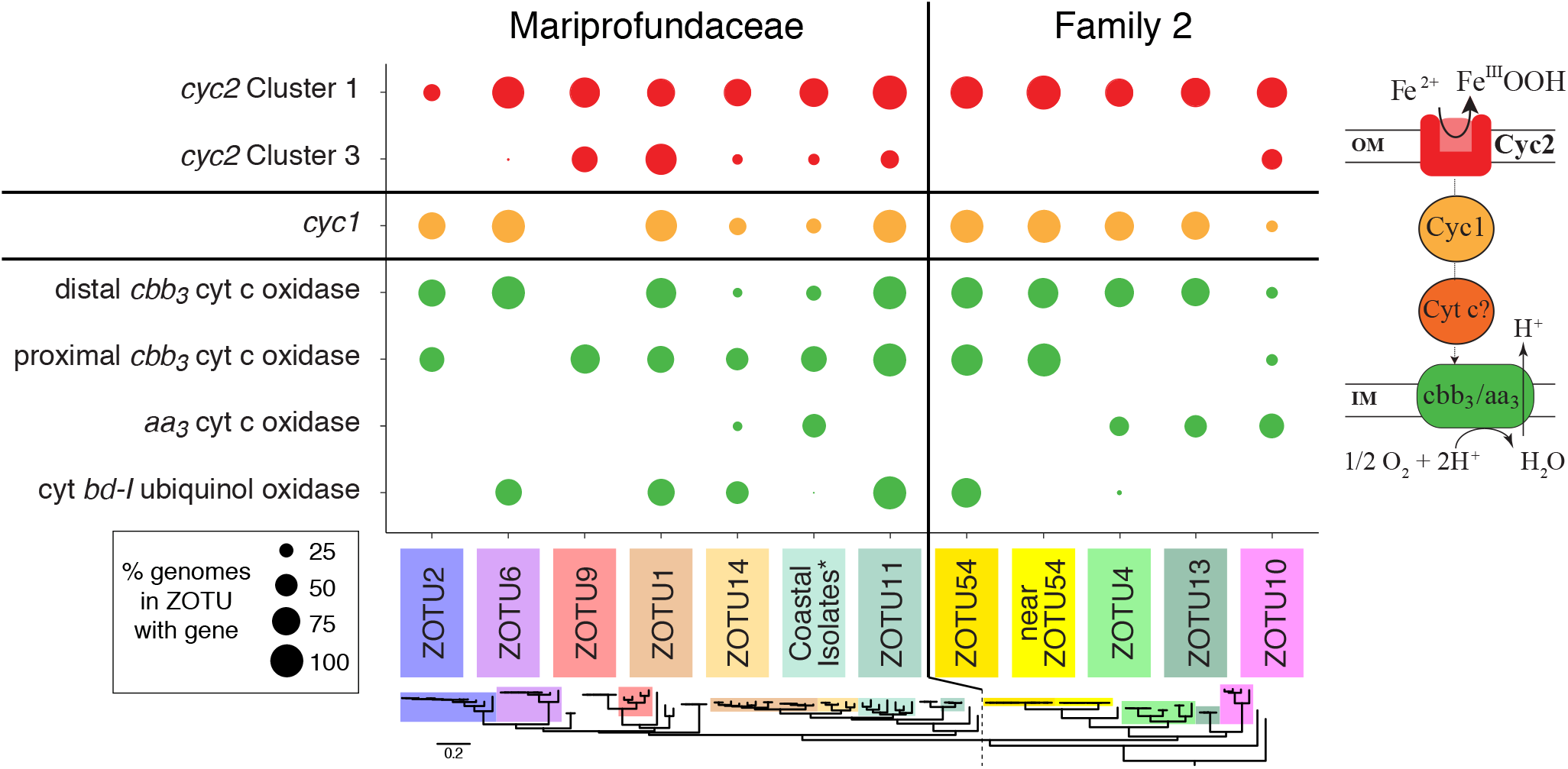
Dot plot showing the distribution of genes from the putative Fe oxidation pathway between major ZOTUs. Each dot represents the % of genomes in the ZOTU which possess the gene of interest. Genes are colored by their relative position within the core putative Fe oxidation pathway, shown to right. ZOTUs are ordered by the reference ribosomal protein tree (bottom), separated into the two families of the Zetaproteobacteria.

In addition to *cyc2*, other proposed genes for the Fe oxidation pathway were also widely distributed in Zetaproteobacteria genomes. Homologs of *cyc1* were present in the genomes of all ZOTUs except ZOTU9 (Figure 4); *cyc1* encodes a di-heme *c*-type cytochrome thought to be a periplasmic electron carrier in the *A. ferrooxidans* Fe oxidation pathway [11]. ZOTU9, which includes the Fe- and H_2_-oxidizing *Ghiorsea* spp. [59], must use another periplasmic electron carrier. Indeed, many other putative periplasmic cytochromes can be found in Zetaproteobacteria genomes (see below). Cyc1 or another electron carrier likely passes electrons to a terminal oxidase or to complex I via complex III (reverse electron transport). Genes for the *bc*_1_ complex were found in all ZOTUs, whereas we found alternative complex III (*ACIII*) genes in only a few Zetaproteobacteria, primarily in ZOTU11 and Family 2 (ZOTUs 4/10/13). We found three types of aerobic terminal oxidases: 1) *cbb_3_*-type cytochrome *c* oxidase, 2) *aa*_3_-type cytochrome *c* oxidase, and 3) cytochrome *bd-I* ubiquinol oxidase. Further, two distinct forms of the *cbb*_3_-type cytochrome *c* oxidase were found, clustering in the proximal and distal *cbb*_3_ subtrees defined by Ducluzeau et al. [60] (**Supplemental Figure 6**). All ZOTUs possess genes for one or more of these terminal oxidases, suggesting that all Zetaproteobacteria are aerobic Fe-oxidizers (Figure 4). Taken together, these findings allow us to update the neutrophilic Fe oxidation pathway model (Figure 5).

**Figure 5.**
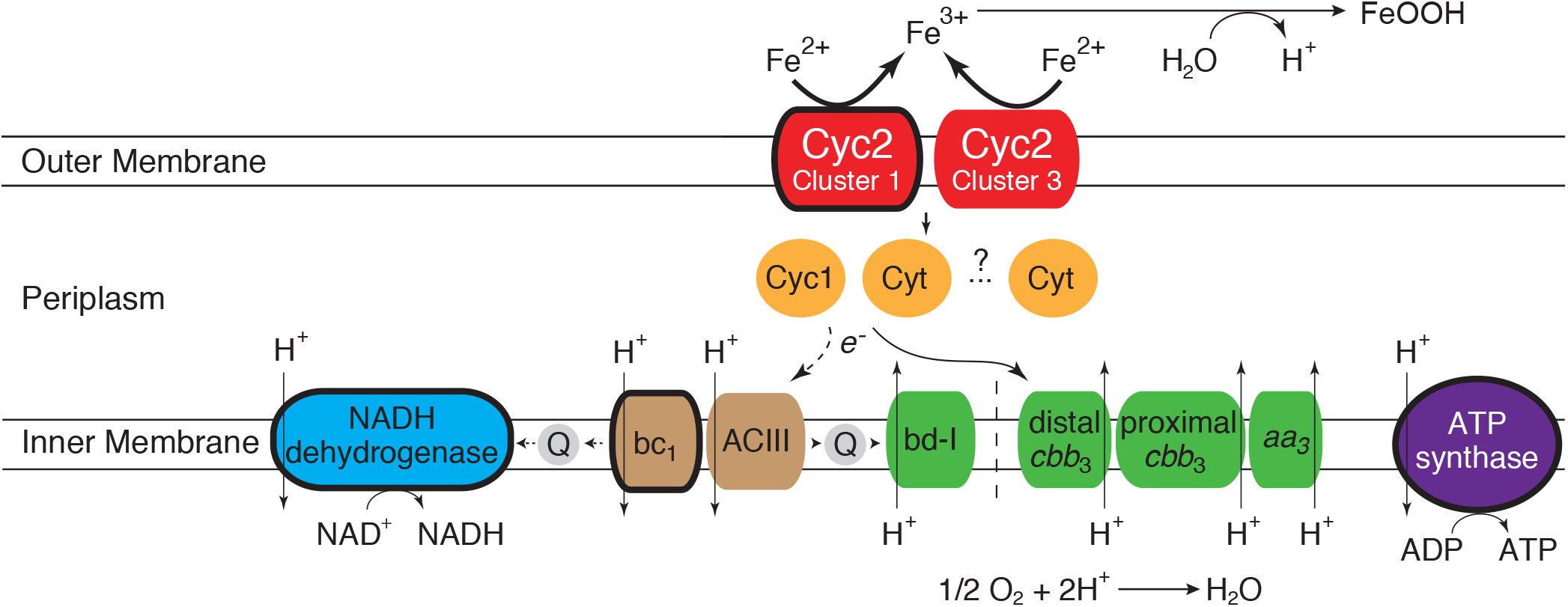
Proposed Fe oxidation pathway model showing variation in the genetic capability of all Zetaproteobacteria. Components that are conserved in all Zetaproteobacteria are outlined with a thick line. Components of the pathway from the same module have the same color.

### *In situ* expression of the putative Fe oxidation pathway

Our next step was to determine whether the putative Fe oxidation pathway genes are expressed in the environment; high expression would lend support for the model. *In situ* expression from six unique ZOTUs in ten different samples (total 21 observations) show that *cyc2* from Cluster 1 are highly expressed in all Zetaproteobacteria and samples, ranging from 3.0x to 555x baseline constitutive gene expression (Table 2). Cluster 1 *cyc2* was frequently the highest expressed gene in the genome, particularly in the Mariprofundaceae (Family 1). Interestingly, *cyc2* expression levels differed between Zetaproteobacteria Families 1 and 2, though expression was still high in all Zetaproteobacteria. The *cyc1* and *cbb_3_*-type terminal oxidase genes are expressed up to 17.3x and 56.6x constitutive gene expression, respectively. On average, this places the *cyc1* and terminal oxidase genes in the 73.4 and 75.4 percentile range in Zetaproteobacteria metatranscriptomic expression, respectively (**Supplemental Figure 7**). This gene expression is consistent with protein expression levels of *M. ferrooxydans* PV-1, where the corresponding proteins were expressed at or above the 87th percentile [14]. In combination, our data suggest that genes in the core model Fe oxidation pathway are highly expressed *in situ* by diverse Zetaproteobacteria under various environmental conditions.

**Table 2.**
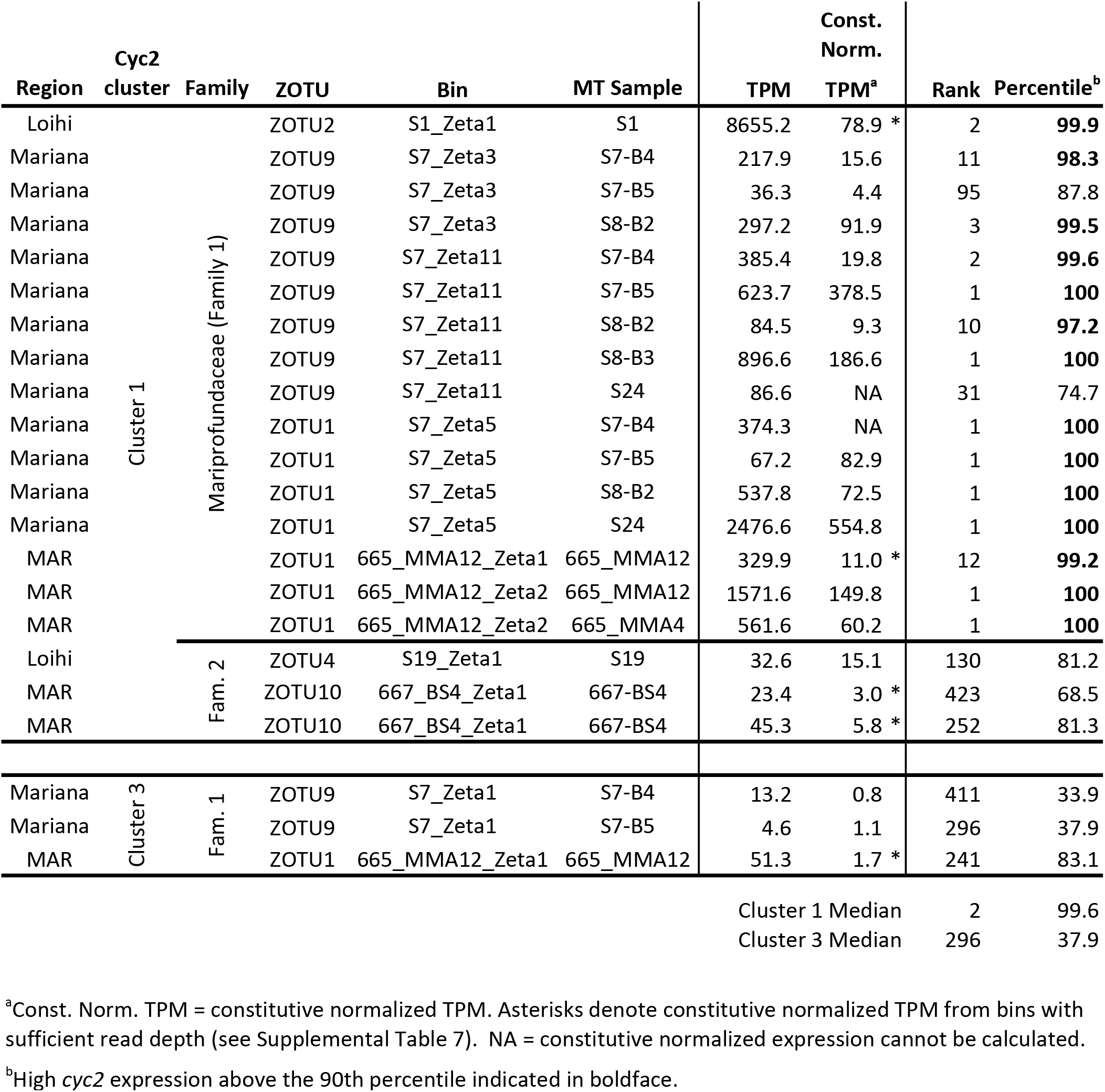
Expression and relative importance of *cyc2* genes from *in situ* samples.

Variation in gene expression may help us resolve which modules are most commonly used during Fe oxidation in the environment. For example, expression of complex III module genes further supports the importance of bc_1_ over ACIII for reverse electron transport. Average expression of *bc*_1_ was 6.6x constitutive expression, while *ACIII* expression was much lower at 0.6x. In ZOTUs with both complexes, *bc*_1_ genes were expressed 1.5x-18.6x higher than the *ACIII* complex. The limited distribution of *ACIII* in only a few Zetaproteobacteria lineages, despite our deep sampling with near complete genomes, combined with its low expression suggests ACIII is not required for Fe oxidation under the sampled conditions.

Comparison of relative *in situ* expression may also help identify genes that may be involved as intermediate electron carriers, particularly in ZOTU9, which lacks *cyc1*. We identified at least 14 different *c*-type cytochrome genes with high expression (>90th percentile) in one or more Zetaproteobacteria genomes (**Supplemental Table 5A**). Some of these genes (*cyc1*, *PC12*, *PC61*, *PC16*, *PC38*) were more highly expressed in some genomes than *cyc2* in the Mariprofundaceae, and all were found in at least one genome where they were more highly expressed than *cyc1*. Interestingly, these putative periplasmic cytochromes were found and expressed at different levels in different Zetaproteobacteria lineages, with some unique to a single ZOTU (e.g. *PC73* in ZOTU2) and most found in several ZOTUs. The cytochromes *c* previously found on a conserved cassette identified in Zetaproteobacteria isolates and SAGs (including *cyc1, PC2*, and *PC3*) [15], are most highly expressed in genomes from ZOTUs 1 and 2. These results help us narrow the list of potential electron carriers in the Zetaproteobacteria.

### Expression of Fe oxidation pathway genes in Fe-amended mat incubations

To link gene expression more specifically with Fe oxidation, we added Fe(II) to mat samples and analyzed metatranscriptomic responses over time. We performed shipboard incubations at Loihi Seamount and Mariana, using fresh Fe mats, live and killed, while monitoring Fe oxidation. Microbes within the mat were actively oxidizing Fe(II) faster than abiotic processes, with the pseudo-first-order Fe oxidation rate constants 3.7x (Loihi) and 5.3x (Mariana) higher in live samples than azide-killed ones (**Supplemental Figure 8**). These results show that we stimulated biotic Fe oxidation, which should lead to increased expression of Fe oxidation genes.

As with the *in situ* samples, *cyc2* was highly expressed in the Zetaproteobacteria during the time series experiments, reaching a maximum of 97.1 to 100 percentile in the four most active Zetaproteobacteria lineages. After Fe(II) was added, there was an increase in total *cyc2* expression (sum of all *cyc2* genes), as well as *cyc2* expression by each individual MAG (Figure 6AB). Expression increased at different rates for each ZOTU, with some peaking earlier and others at the end of the experiment. Expression of *cyc2* increased less for the most abundant Zetaproteobacteria (e.g. S6_Zeta1), which already had high expression of *cyc2* prior to Fe(II) amendment. Less abundant Zetaproteobacteria (S6_Zeta11/S6_Zeta23) were also expressing *cyc2* prior to Fe(II) amendment, but had a much larger change in expression (5.7-6.5x), reached their maximum quickly, and maintained a higher expression over the course of the experiment. Overall, while the degree and timing of response differed, all Zetaproteobacteria increased their *cyc2* expression in response to Fe(II) amendment.

**Figure 6.**
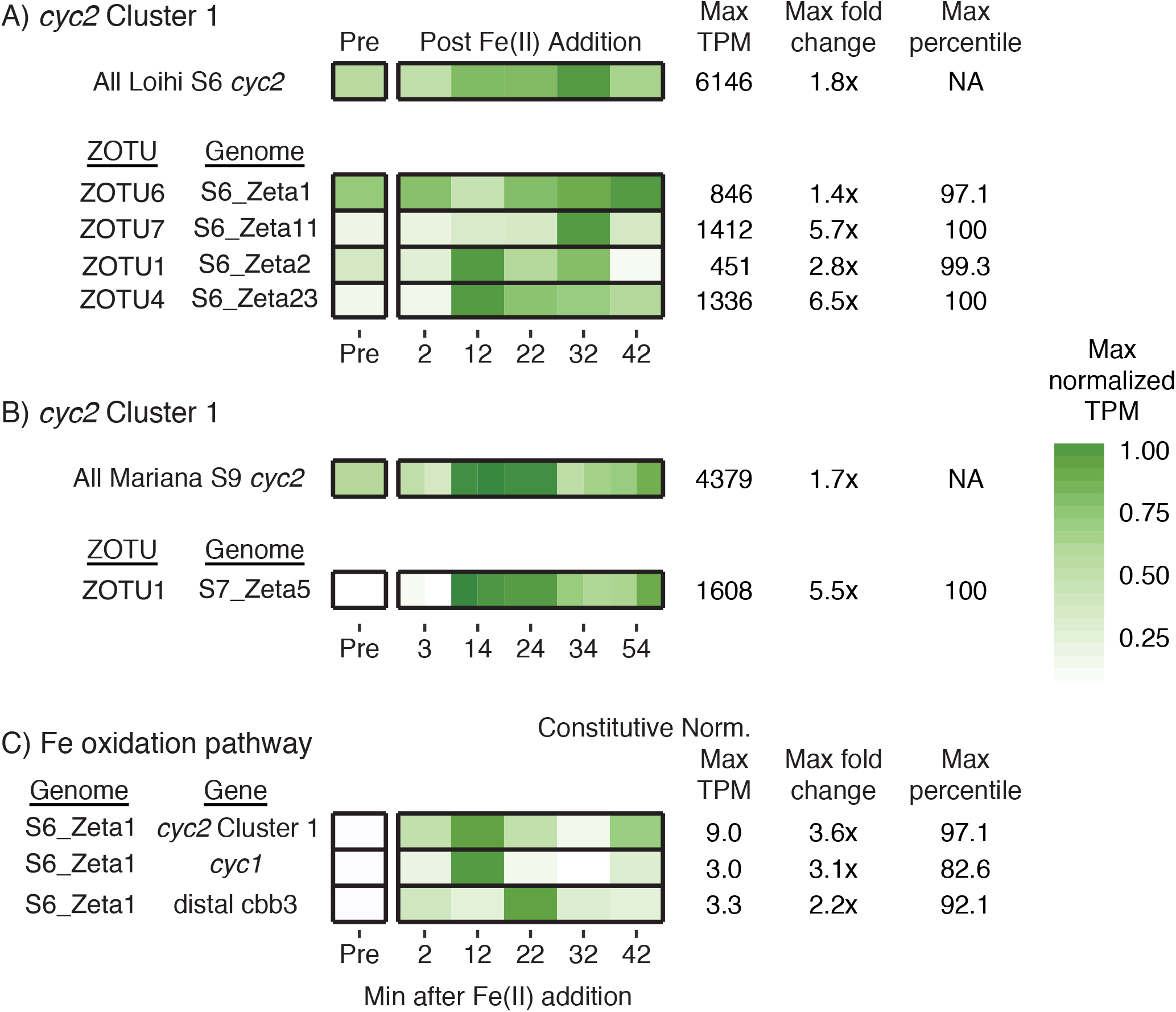
Normalized TPM expression changes for *cyc2* and other Fe oxidation pathway genes in the Fe(II) amendment experiments, showing increases post Fe(II) addition. (A-B) TPM expression changes are shown for all Cluster 1 *cyc2* and for *cyc2* from specific Zetaproteobacteria MAGs at (A) Loihi and (B) Mariana (duplicates shown). (C) Constitutive-normalized expression changes shown for the Fe oxidation pathway in MAG S6_Zeta1. S9 MT data mapped to the S7 MG for expression estimates of MAGs in (B). TPM values were max-normalized to emphasize peak expression.

Other genes in the Fe oxidation pathway also generally followed this trend, with *cyc1* expression in 4 of 5 genomes and *cbb*_3_–type terminal oxidase genes in 2 of 4 genomes also increasing after Fe(II) addition. However, low read recruitment depth led to substantial noise. To correct for this noise, we normalized expression using six constitutively expressed genes, and focused on expression patterns in S6_Zeta1, which had high read depth (Figure 6C). Constitutive normalized expression of S6_Zeta1 shows a similar pattern of *cyc2* expression change over the time series compared to expression patterns prior to constitutive normalization, with a maximum increase of 3.6x after Fe(II) amendment. The genes encoding Cyc1 and the *cbb*_3_-type terminal oxidase also increased after Fe(II) addition, reaching a maximum fold change of 1.5 - 3x. This trend was also observed for 7 of 8 putative periplasmic cytochromes more highly expressed than *cyc1*, increasing 2.1-4.5x over the course of the experiment (**Supplemental Table 5B**). These results suggest that the Fe(II) amendment increased the expression of many genes thought to be in the Fe oxidation pathway.

If the Zetaproteobacteria represented by the S6_Zeta1 genome is an autotrophic Fe-oxidizer, Fe(II) amendment should stimulate genes for carbon fixation, central metabolism, and growth. Like *cyc2* expression, genes for central metabolic pathways, including the TCA cycle, increased in expression 2.0-3.4x over one hour in S6_Zeta1 from ZOTU6 (**Supplemental Figure 9**). Similarly, expression of genes for glycogen synthesis increased 1.8x in the first two minutes after Fe(II) addition. Carbon fixation genes increased 1.9x in the first 12 minutes. Some of the highest fold changes after Fe(II) amendment were seen in genes related to proper protein folding (molecular chaperones *groEL, groES*, and *dnaK*) and membrane protein quality control (*htpX-*type protease) [61]. For example, *groEL* increased 119x after Fe(II) addition. Though these gene responses may correspond with shock to the cell after Fe(II) amendment, these genes were also highly expressed under *in situ* conditions in S1_Zeta1, which suggests that they may be necessary for promoting active Fe oxidation in the environment. Together, these data suggest Fe(II) amendment stimulated genes for both Fe oxidation and growth.

## DISCUSSION

The main objective of this study was to assess the Cyc2-based Fe oxidation pathway in neutrophilic FeOB, using environmental metagenomics and metatranscriptomics of marine Fe mats. This work contributes 53 new near-complete Zetaproteobacteria genomes paired with expression profiles *in situ* and from incubations, along with a comprehensive Cyc2 phylogeny. Using these, we have characterized the distribution and usage of the model Fe oxidation pathway across the full range of known Zetaproteobacteria in Fe mats at three geographically-distinct venting regions. The emerging pattern shows that the pathway as a whole is highly expressed, with increased expression in all Fe pathway genes following Fe(II) amendment. The *cyc2* gene is among the highest expressed and is the only gene in the pathway shared by all Zetaproteobacteria, suggesting it plays a key role in Fe oxidation.

### Assessing the Fe oxidation pathway model through Zetaproteobacteria comparative genomics

In this study, we assessed the current model for Fe oxidation by comparing genomes representing the full diversity of the Zetaproteobacteria (Figure 1B). Since the Zetaproteobacteria are thought to be an entire class of Fe-oxidizing bacteria, all genomes should have an Fe-oxidizing pathway, which may be conserved. Our results show that overall, the basic model of neutrophilic Fe oxidation in the Zetaproteobacteria holds (Figure 4). All Zetaproteobacteria lineages, including novel ones presented in this study, possess genes encoding the putative Fe oxidase Cyc2, an intermediate electron carrier, and a terminal oxidase. However, our survey shows that each of these components can have multiple versions, suggesting that the pathway contains interchangeable modules, as depicted in our updated model (Figure 5). Across all Zetaproteobacteria, there are two types of Cyc2, multiple potential periplasmic cytochrome electron carriers, and four terminal oxidases. The variations are likely linked with specific adaptations related to niche, with some genomes possessing multiple versions of certain components, perhaps to span multiple niches [62]. Within each ZOTU, individual genomes possessed pathway gene variations consistent with the ZOTU as a whole, even though some genomes were missing genes that a majority of others in the ZOTU possessed. These false negatives could result from incomplete MAG genomes, which is why we focused on ZOTUs. Accounting for modularity and genome variability within ZOTUs, comparative genomics has confirmed that this Fe oxidation pathway is common to all Zetaproteobacteria lineages.

### Support for *cyc2* as an Fe oxidation gene in neutrophilic Fe-oxidizers

Cyc2 homologs in neutrophilic FeOB are commonly referred to as “putative Fe oxidases” based on homology to the functionally-characterized *A. ferrooxidans* Cyc2, though sequence similarity is low. Indeed, our phylogenetic analysis shows that Cyc2 sequences are highly diverse, and most of the neutrophilic FeOB Cyc2 homologs fall into Cluster 1 (Figure 3). This cluster forms a distinct group from the clusters containing biochemically characterized Fe oxidases: Cyc2 from *A. ferrooxidans* (Cluster 2) and Cyt_572_ from *Leptospirillum* (Cluster 3). Although none of the Cluster 1 Cyc2 have been biochemically characterized, the high bootstrap support suggests a common function, and the prevalence of sequences from Fe-oxidizing isolates strongly suggests involvement in Fe oxidation.

Another clue to function lies in the expression of *cyc2* in Fe mat environments, where Fe oxidation by the Zetaproteobacteria is required for carbon fixation and growth. Genes central to fitness are often highly expressed [63, 64]. We measured *cyc2* expression in five different active Fe mats from three different hydrothermal vent fields, and confirmed that *cyc2* is frequently the highest expressed gene. This is true in diverse Zetaproteobacteria lineages (Table 2), suggesting that the pathway is important to Zetaproteobacteria fitness in many different environments. The level of *cyc2* expression in the Zetaproteobacteria is consistent with the high expression in Fe-oxidizing isolates of *Acidithiobacillus* sp. (often above microarray detection limits) and *Ferrovum* sp. (8x above average) [65–67]. In the environment, the neutrophilic Gallionellaceae have been shown to express *cyc2* highly, up to the 100th percentile in an Fe-rich aquifer [68]. Together with the expression of *cyc1* and terminal oxidase genes, this shows that the putative Fe oxidation pathway is consistently expressed under Fe-oxidizing conditions. The especially high expression of *cyc2* across various Fe-oxidizing taxa and Fe-oxidizing environments supports its role in Fe oxidation in both acidophiles and neutrophiles.

To link *cyc2* to Fe oxidation, we followed its expression when Fe(II) was added to Fe mat samples. In separate experiments at Loihi and Mariana, Fe(II) amendment resulted in both active biotic Fe oxidation (**Supplemental Figure 8**) and increased *cyc2* gene expression. This increase in expression was found not only for the whole sample, but also in every Zetaproteobacteria genome detected within these samples (Figure 6). Although there was variation in the timing and magnitude of the response, which may be lineage specific, the fact that expression increased in all Zetaproteobacteria suggests that *cyc2* expression is stimulated by the presence of Fe(II). The Fe(II) amendment also resulted in increases in carbon fixation and central metabolism gene expression, suggesting a link between *cyc2* expression, neutrophilic Fe oxidation, and growth (**Supplemental Figure 9**).

### Can the *cyc2* gene be used as a marker for Fe oxidation?

Unlike many other energy metabolisms, neutrophilic Fe oxidation is challenging to track in the environment due to lack of an isotopic signature and difficulties distinguishing biotic from abiotic Fe oxides. Until recently, there have not been any candidates for a widely-applicable Fe oxidation genetic marker; instead, it seemed that there were many different potential Fe oxidases, with varying levels of functional verification (e.g. [6, 7, 69]). Our work adds to the mounting evidence that Cyc2 is an Fe oxidase. The *cyc2* gene is widely distributed across many Fe-oxidizing lineages, with homologs in acidophilies and neutrophiles. Specifically for neutrophiles, *cyc2* is common across the well-studied neutrophilic chemolithotrophs Gallionellaceae and Zetaproteobacteria. As we have sequenced more of these neutrophilic FeOB genomes, this association has held true [9, 13, 15, 21, 70]. However, our Cyc2 phylogeny has identified a substantial number of organisms that have not yet been shown or tested to be capable of Fe oxidation, work that could bolster confidence. In all, the *cyc2* gene is a promising genomic marker of the capacity for Fe oxidation across many different Fe-oxidizing lineages, including neutrophiles.

Not only is *cyc2* common to all well-established neutrophilic Fe-oxidizers, it is also highly expressed in environments where neutrophilic Fe-oxidizers predominate (this study; [68]). This opens the possibility of *cyc2* expression levels as an indicator of microbial Fe oxidation activity. Indeed, when we stimulated Zetaproteobacteria Fe oxidation in incubations, expression of *cyc2* increased along with an increase in carbon fixation and central metabolism genes. This is consistent with Fe oxidation-fueled chemolithoautotrophic growth, and so relative *cyc2* expression levels can correspond to increases in Fe oxidation activity. However, our results suggest that *cyc2* expression levels may not be easily related to Fe oxidation activity in the environment. All Zetaproteobacteria in our samples were expressing *cyc2* prior to Fe(II) amendment, when there was no detectable dissolved Fe(II). This could represent baseline expression by obligate Fe-oxidizers, which always need to be prepared for Fe oxidation. In this case relative changes in *cyc2* expression would remain more informative for activity. Alternatively, *cyc2* expression before Fe(II) amendment could result from cryptic cycling of Fe between Fe-oxidizers and reducers [71]. Such cryptic cycling would make developing a genetic marker for activity even more important for tracking Fe oxidation activity. Because of these potential complications, further transcriptomics experiments should focus on isolates or microcosms without Fe-reducers. In combination with our results, such experiments will help us understand how to use *cyc2* expression levels to interpret Fe oxidation activity in the environment.

### Conclusions

Using paired metagenomes and metatranscriptomes from the Zetaproteobacteria, we have been able to demonstrate that the Cyc2-based neutrophilic Fe oxidation pathway is widespread and highly expressed in the environment, validating the environmental importance of the pathway. We have shown that the Cluster 1 *cyc2* gene, conserved in the Zetaproteobacteria and other neutrophilic Fe-oxidizers, is highly expressed in multiple Fe mat environments and is stimulated by Fe(II) addition, suggesting it may be regulated. This makes *cyc2* an excellent marker of Fe oxidation capability, and may allow us to detect and monitor the activity of Fe-oxidizers in the environment. However, to correlate expression with activity, further efforts should focus on testing the regulation of *cyc2* in diverse organisms and simple communities. The phylogeny of Cyc2 shows at least three distinct clusters, with some neutrophilic Fe-oxidizers possessing multiple copies (e.g. Clusters 1 and 3 in the Zetaproteobacteria). It is unclear why there are multiple *cyc2* in a single genome, though organisms may use different Fe oxidases under different kinds of Fe-oxidizing conditions. If so, it would be valuable to monitor each form of *cyc2* individually. Without a marker of activity, the roles of neutrophilic Fe-oxidizers have been virtually invisible outside of model Fe-oxidizing environments, like Fe microbial mats. By applying our findings to other environments, we can start to reveal how Fe-oxidizing microbes drive key biogeochemical cycles in the varied marine and freshwater habitats where they thrive.

## Acknowledgements

SMM and CSC drafted the manuscript. All authors contributed to experimental design and editing. DAB, BTG, and SMM performed geochemical analysis. SMM, JBS, BTG, and CSC implemented shipboard Fe(II) amendment experiments. SMM, SWP, and CSC conducted bioinformatics analysis. We acknowledge George W. Luther, III, for generous ship time and geochemical data from the MAR. We thank the captains and crew of the R/Vs Knorr, Thompson, and Revelle and ROV Jason II. We also acknowledge Anna Leavitt, Arne Sturm, Angelos Hannides, and Karyn Rogers for their assistance with the shipboard experiments; Kevin Roe for chemical analyses of fluid samples; Vadesse Noundou for assistance with Zetaproteobacteria central metabolism; Ryan Moore and Karol Miaskiewicz for bioinformatics assistance; Jennifer Biddle, David Emerson, Thomas Hanson, and Jessica Keffer for their comments on this manuscript. We also thank the University of Delaware Sequencing and Genotyping Center for their help with sample preparation and sequencing, in particular Bruce Kingham, Summer Thompson, and Olga Shevchenko. This work was funded by NSF OCE-1155290 and ONR N00014-17-1-2640 (to CSC), C-DEBI (contribution ####) (to JBS), NSF OCE-1061827 and OCE-1031947 (to BTG), NOAA/PMEL (contribution ####), Ocean Exploration and Research, and JISAO (contribution #2019-xxx) (to DAB), NSF EAR-1833525 (to CSC and SWP), and NIGMS P20 GM103446 (to SWP), in addition to two Delaware Space Grant Fellowships (NASA Grant NNX10AN63H) and the University of Delaware Dissertation Fellowship to SMM. Computational infrastructure support by the University of Delaware CBCB Core Facility funded by Delaware INBRE (NIH NIGMS P20 GM103446) and the Delaware Biotechnology Institute.

## References

1. Emerson D, Fleming EJ, McBeth JM. Iron-oxidizing bacteria: an environmental and genomic perspective. Annu Rev Microbiol 2010; 64: 561–583.

2. Chan CS, Fakra SC, Emerson D, Fleming EJ, Edwards KJ. Lithotrophic iron-oxidizing bacteria produce organic stalks to control mineral growth: implications for biosignature formation. ISME J 2011; 5: 717–727.

3. Laufer K, Nordhoff M, Halama M, Martinez RE, Obst M, Nowak M, et al. Microaerophilic Fe(II)-oxidizing Zetaproteobacteria isolated from low-Fe marine coastal sediments: Physiology and characterization of their twisted stalks. Appl Environ Microbiol 2017; 83: e03118–16.

4. Kendall B, Anbar AD, Kappler A, Konhauser KO. The global iron cycle. In: Knoll AH, Canfield DE, Konhauser KO (eds). Fundamentals of Geobiology, 1st ed. 2012. Blackwell Publishing Ltd., pp 65–92.

5. Liu J, Wang Z, Belchik SM, Edwards MJ, Liu C, Kennedy DW, et al. Identification and characterization of MtoA: a decaheme c-type cytochrome of the neutrophilic Fe(II)-oxidizing bacterium Sideroxydans lithotrophicus ES-1. Front Microbiol 2012; 3: 37.

6. White GF, Edwards MJ, Gomez-Perez L, Richardson DJ, Butt JN, Clarke TA. Mechanisms of bacterial extracellular electron exchange. Adv Microb Physiol 2016; 68: 87–138.

7. He S, Barco RA, Emerson D, Roden EE. Comparative genomic analysis of neutrophilic iron(II) oxidizer genomes for candidate genes in extracellular electron transfer. Front Microbiol 2017; 8: 1584.

8. Kato S, Ohkuma M, Powell DH, Krepski ST, Oshima K, Hattori M, et al. Comparative genomic insights into ecophysiology of neutrophilic, microaerophilic iron oxidizing bacteria. Front Microbiol 2015; 6: 1265.

9. Fullerton H, Hager KW, McAllister SM, Moyer CL. Hidden diversity revealed by genome-resolved metagenomics of iron-oxidizing microbial mats from Lo’ihi Seamount, Hawai’i. ISME J 2017; 11: 1900–1914.

10. McAllister SM, Moore RM, Gartman A, Luther GW, Emerson D, Chan CS. The Fe(II)-oxidizing Zetaproteobacteria: Historical, ecological, and genomic perspectives. FEMS Microbiol Ecol 2019; fiz015.

11. Castelle C, Guiral M, Malarte G, Ledgham F, Leroy G, Brugna M, et al. A new iron-oxidizing/O_2_-reducing supercomplex spanning both inner and outer membranes, isolated from the extreme acidophile Acidithiobacillus ferrooxidans. J Biol Chem 2008; 283: 25803–25811.

12. Jeans C, Singer SW, Chan CS, VerBerkmoes NC, Shah M, Hettich RL, et al. Cytochrome 572 is a conspicuous membrane protein with iron oxidation activity purified directly from a natural acidophilic microbial community. ISMEJ 2008; 2: 542–550.

13. Emerson D, Field EK, Chertkov O, Davenport KW, Goodwin L, Munk C, et al. Comparative genomics of freshwater Fe-oxidizing bacteria: implications for physiology, ecology, and systematics. Front Microbiol 2013; 4: 254.

14. Barco RA, Emerson D, Sylvan JB, Orcutt BN, Jacobson Meyers ME, Ramírez GA, et al. New insight into microbial iron oxidation as revealed by the proteomic profile of an obligate iron-oxidizing chemolithoautotroph. Appl Environ Microbiol 2015; 81: 5927–5937.

15. Field EK, Sczyrba A, Lyman AE, Harris CC, Woyke T, Stepanauskas R, et al. Genomic insights into the uncultivated marine Zetaproteobacteria at Loihi Seamount. ISME J 2015; 9: 857–870.

16. Emerson D, Rentz JA, Lilburn TG, Davis RE, Aldrich H, Chan C, et al. A novel lineage of proteobacteria involved in formation of marine Fe-oxidizing microbial mat communities. PLoS One 2007; 2: e667.

17. Makita H, Kikuchi S, Mitsunobu S, Takaki Y, Yamanaka T, Toki T, et al. Comparative analysis of microbial communities in iron-dominated flocculent mats in deep-sea hydrothermal environments. Appl Environ Microbiol 2016; 82: 5741–5755.

18. Barco RA, Hoffman CL, Ramírez GA, Toner BM, Edwards KJ, Sylvan JB. In-situ incubation of iron-sulfur mineral reveals a diverse chemolithoautotrophic community and a new biogeochemical role for Thiomicrospira. Environ Microbiol 2017; 19: 1322–1337.

19. Probst AJ, Castelle CJ, Singh A, Brown CT, Anantharaman K, Sharon I, et al. Genomic resolution of a cold subsurface aquifer community provides metabolic insights for novel microbes adapted to high CO_2_ concentrations. Environ Microbiol 2017; 19: 459–474.

20. Meyer JL, Jaekel U, Tully BJ, Glazer BT, Wheat CG, Lin H-T, et al. A distinct and active bacterial community in cold oxygenated fluids circulating beneath the western flank of the Mid-Atlantic ridge. Sci Rep 2016; 6: 22541.

21. Garrison CE, Price KA, Field EK. Environmental evidence for and genomic insight into the preference of iron-oxidizing Bacteria for more-corrosion-resistant stainless steel at higher salinities. Appl Environ Microbiol 2019; 85: e00483–19.

22. McAllister SM, Davis RE, McBeth JM, Tebo BM, Emerson D, Moyer CL. Biodiversity and emerging biogeography of the neutrophilic iron-oxidizing Zetaproteobacteria. Appl Environ Microbiol 2011; 77: 5445–5457.

23. Hager KW, Fullerton H, Butterfield DA, Moyer CL. Community structure of lithotrophically-driven hydrothermal microbial mats from the Mariana Arc and Back-Arc. Front Microbiol 2017; 8: 1578.

24. Vander Roost J, Thorseth IH, Dahle H. Microbial analysis of Zetaproteobacteria and co-colonizers of iron mats in the Troll Wall Vent Field, Arctic Mid-Ocean Ridge. PLoS One 2017; 12: e0185008.

25. Vander Roost J, Daae FL, Steen IH, Thorseth IH, Dahle H. Distribution patterns of iron-oxidizing Zeta- and Beta-Proteobacteria from different environmental settings at the Jan Mayen Vent Fields. Front Microbiol 2018; 9: 3008.

26. Chan CS, McAllister SM, Leavitt AH, Glazer BT, Krepski ST, Emerson D. The architecture of iron microbial mats reflects the adaptation of chemolithotrophic iron oxidation in freshwater and marine environments. Front Microbiol 2016; 7: 796.

27. Breier JA, Gomez-Ibanez D, Reddington E, Huber JA, Emerson D. A precision multi-sampler for deep-sea hydrothermal microbial mat studies. Deep Sea Res Part I Oceanogr Res Pap 2012; 70: 83–90.

28. MacDonald DJ, Findlay AJ, McAllister SM, Barnett JM, Hredzak-Showalter P, Krepski ST, et al. Using in situ voltammetry as a tool to identify and characterize habitats of iron-oxidizing bacteria: from fresh water wetlands to hydrothermal vent sites. Environ Sci Process Impacts 2014; 16: 2117–2126.

29. Glazer BT, Rouxel OJ. Redox speciation and distribution within diverse iron-dominated microbial habitats at Loihi Seamount. Geomicrobiol J 2009; 26: 606–622.

30. Butterfield DA, Roe KK, Lilley MD, Huber JA, Baross JA, Embley RW, et al. Mixing, reaction and microbial activity in the sub-seafloor revealed by temporal and spatial variation in diffuse flow vents at Axial Volcano. The Subseafloor Biosphere at Mid-Ocean Ridges. 2013. American Geophysical Union (AGU).

31. Stookey LL. Ferrozine--a new spectrophotometric reagent for iron. Anal Chem 1970; 42: 779–781.

32. Klueglein N, Kappler A. Abiotic oxidation of Fe(II) by reactive nitrogen species in cultures of the nitrate-reducing Fe(II) oxidizer Acidovorax sp. BoFeN1 - questioning the existence of enzymatic Fe(II) oxidation. Geobiology 2013; 11: 180–190.

33. McAllister SM, Moore RM, Chan CS. ZetaHunter, a reproducible taxonomic classification tool for tracking the ecology of the Zetaproteobacteria and other poorly resolved taxa. Microbiol Resour Announc 2018; 7: e00932–18.

34. Bolger AM, Lohse M, Usadel B. Trimmomatic: A flexible trimmer for Illumina sequence data. Bioinformatics 2014; 30: 2114–2120.

35. Magoč T, Salzberg SL. FLASH: fast length adjustment of short reads to improve genome assemblies. Bioinformatics 2011; 27: 2957–2963.

36. Nurk S, Meleshko D, Korobeynikov A, Pevzner PA. metaSPAdes: a new versatile metagenomic assembler. Genome Res 2017; 27: 824–834.

37. Wu Y-W, Tang Y-H, Tringe SG, Simmons BA, Singer SW. Max Bin: an automated binning method to recover individual genomes from metagenomes using an expectation-maximization algorithm. Microbiome 2014; 2: 26.

38. Kang DD, Froula J, Egan R, Wang Z. MetaBAT, an efficient tool for accurately reconstructing single genomes from complex microbial communities. PeerJ 2015; 3: e1165.

39. Alneberg J, Bjarnason BS, De Bruijn I, Schirmer M, Quick J, Ijaz UZ, et al. Binning metagenomic contigs by coverage and composition. Nat Methods 2014; 11: 1144–1146.

40. Graham ED, Heidelberg JF, Tully BJ. BinSanity: unsupervised clustering of environmental microbial assemblies using coverage and affinity propagation. PeerJ 2017; 5: e3035.

41. Sieber CMK, Probst AJ, Sharrar A, Thomas BC, Hess M, Tringe SG, et al. Recovery of genomes from metagenomes via a dereplication, aggregation and scoring strategy. Nat Microbiol 2018; 3: 836–843.

42. Eren AM, Esen ÖC, Quince C, Vineis JH, Morrison HG, Sogin ML, et al. Anvi’o: an advanced analysis and visualization platform for ‘omics data. PeerJ 2015; 3: e1319.

43. Parks DH, Imelfort M, Skennerton CT, Hugenholtz P, Tyson GW. CheckM: assessing the quality of microbial genomes recovered from. Genome Res 2015; 25: 1043–1055.

44. Darling AE, Jospin G, Lowe E, Matsen FA, Bik HM, Eisen JA. Phylo Sift: phylogenetic analysis of genomes and metagenomes. PeerJ 2014; 2: e243.

45. Overbeek R, Olson R, Pusch GD, Olsen GJ, Davis JJ, Disz T, et al. The SEED and the Rapid Annotation of microbial genomes using Subsystems Technology (RAST). Nucleic Acids Res 2014; 42: D206–D214.

46. Galperin MY, Makarova KS, Wolf YI, Koonin E V. Expanded microbial genome coverage and improved protein family annotation in the COG database. Nucleic Acids Res 2015; 43: D261–D269.

47. Kanehisa M, Sato Y, Morishima K. BlastKOALA and GhostKOALA: KEGG Tools for functional characterization of genome and metagenome sequences. J Mol Biol 2016; 428: 726–731.

48. Camacho C, Coulouris G, Avagyan V, Ma N, Papadopoulos J, Bealer K, et al. BLAST+: Architecture and applications. BMC Bioinformatics 2009; 10: 421.

49. Stamatakis A, Hoover P, Rougemont J. A rapid bootstrap algorithm for the RAxML web servers. Syst Biol 2008; 57: 758–71.

50. Chen I-MA, Markowitz VM, Chu K, Palaniappan K, Szeto E, Pillay M, et al. IMG/M: Integrated genome and metagenome comparative data analysis system. Nucleic Acids Res 2017; 45: D507–D516.

51. Benson DA, Cavanaugh M, Clark K, Karsch-Mizrachi I, Lipman DJ, Ostell J, et al. GenBank. Nucleic Acids Res 2013; 41: D37–D42.

52. Kopylova E, Noé L, Touzet H. SortMeRNA: Fast and accurate filtering of ribosomal RNAs in metatranscriptomic data. Bioinformatics 2012; 28: 3211–3217.

53. Langmead B, Salzberg SL. Fast gapped-read alignment with Bowtie 2. Nat Methods 2012; 9: 357–359.

54. Quinlan AR, Hall IM. BEDTools: A flexible suite of utilities for comparing genomic features. Bioinformatics 2010; 26: 841–842.

55. Wagner GP, Kin K, Lynch VJ. Measurement of mRNA abundance using RNA-seq data: RPKM measure is inconsistent among samples. Theory Biosci 2012; 131: 281–285.

56. Rocha DJP, Santos CS, Pacheco LGC. Bacterial reference genes for gene expression studies by RT-qPCR: survey and analysis. Antonie Van Leeuwenhoek 2015; 108: 685–693.

57. Scott JJ, Glazer BT, Emerson D. Bringing microbial diversity into focus: high-resolution analysis of iron mats from the Lō’ihi Seamount. Environ Microbiol 2017; 19: 301–316.

58. Scott JJ, Breier JA, Luther, III GW, Emerson D. Microbial iron mats at the Mid-Atlantic Ridge and evidence that Zetaproteobacteria may be restricted to iron-oxidizing marine systems. PLoS One 2015; 10: e0119284.

59. Mori JF, Scott JJ, Hager KW, Moyer CL, Küsel K, Emerson D. Physiological and ecological implications of an iron- or hydrogen-oxidizing member of the Zetaproteobacteria, *Ghiorsea bivora*, gen. nov., sp. nov. ISME J 2017; 11: 2624–2636.

60. Ducluzeau AL, Ouchane S, Nitschke W. The cbb_3_ oxidases are an ancient innovation of the domain Bacteria. Mol Biol Evol 2008; 25: 1158–1166.

61. Sakoh M, Ito K, Akiyama Y. Proteolytic activity of HtpX, a membrane-bound and stress-controlled protease from *Escherichia coli*. J Biol Chem 2005; 280: 33305–33310.

62. McAllister SM. The Zetaproteobacteria: Ecology and metabolic functions of a model neutrophilic Fe-oxidizing clade. University of Delaware Dissertation. 2019.

63. Frias-Lopez J, Shi Y, Tyson GW, Coleman ML, Schuster SC, Chisholm SW, et al. Microbial community gene expression in ocean surface waters. PNAS 2008; 105: 3805–3810.

64. Urich T, Lanzén A, Stokke R, Pedersen RB, Bayer C, Thorseth IH, et al. Microbial community structure and functioning in marine sediments associated with diffuse hydrothermal venting assessed by integrated meta-omics. Environ Microbiol 2014; 16: 2699–2710.

65. Yarzábal A, Appia-Ayme C, Ratouchniak J, Bonnefoy V. Regulation of the expression of the Acidithiobacillus ferrooxidans *rus* operon encoding two cytochromes *c*, a cytochrome oxidase and rusticyanin. Microbiology 2004; 150: 2113–2123.

66. Quatrini R, Appia-Ayme C, Denis Y, Ratouchniak J, Veloso F, Valdes J, et al. Insights into the iron and sulfur energetic metabolism of *Acidithiobacillus ferrooxidans* by microarray transcriptome profiling. Hydrometallurgy 2006; 83: 263–272.

67. Ullrich SR, Poehlein A, Levicán G, Mühling M, Schlömann M. Iron targeted transcriptome study draws attention to novel redox protein candidates involved in ferrous iron oxidation in “Ferrovum” sp. JA12. Res Microbiol 2018; 169: 618–627.

68. Jewell TNM, Karaoz U, Brodie EL, Williams KH, Beller HR. Metatranscriptomic evidence of pervasive and diverse chemolithoautotrophy relevant to C, S, N, and Fe cycling in a shallow alluvial aquifer. ISME J 2016; 10: 2106–2117.

69. Ilbert M, Bonnefoy V. Insight into the evolution of the iron oxidation pathways. Biochim Biophys Acta 2013; 1827: 161–75.

70. Crowe SA, Hahn AS, Morgan-Lang C, Thompson KJ, Simister RL, Llirós M, et al. Draft genome sequence of the pelagic photoferrotroph Chlorobium phaeoferrooxidans. Genome Announc 2017; 5: e01584–16.

71. Emerson D. Potential for iron-reduction and iron-cycling in iron oxyhydroxide-rich microbial mats at Loihi Seamount. Geomicrobiol J 2009; 26: 639–647.

