## Supplemental Material for "Validating the Cyc2 neutrophilic Fe oxidation pathway using meta-omics of Zetaproteobacteria iron mats at marine hydrothermal vents"


**PDF document includes:**


- **Supplemental methods (Pages 2-5)**
- **Supplemental text (Pages 5-6)**
- **Supplemental references (Pages 6-9)**
- **7 Supplemental Figures**


**Supplemental Tables (xlsx):**

**Supplemental Table 1:** Sample names, origin, and type for this study.
**Supplemental Table 2:** Metagenome and metatranscriptome sequencing information and MAG recovery results.
**Supplemental Table 3:** Statistics for all bins from this study, including our own samples and reassembled data from Fullerton et al. (2017).
**Supplemental Table 4:** Zetaproteobacteria genomes used in comparative genomics, concatenated ribosomal protein phylogenetic tree, and gene expression estimates.
**Supplemental Table 5:** (A) Percentile expression for each putative periplasmic cytochrome (PC) compared to cyc2. (B) Constitutive normalized expression for PC compared with cyc2 in genome S6_Zeta1.
**Supplemental Table 6:** Statistics for the pTXB1 internal RNA standard.
**Supplemental Table 7:** Average constitutive gene expression (TPM) for each unique MT mapped against the Zetaproteobacteria bins.

### Supplemental Methods

#### RNA extraction
RNA samples were extracted using the NucleoSpin RNA kit (Macherey-Nagel, Bethlehem, PA, USA), with modifications from the manufacturer's instructions: 1) RNase inhibitor SUPERase-In (Invitrogen, Carlsbad, CA, USA) was added prior to lysis (300 U), prior to loading on the membrane filter (300 U), and prior to elution (100 U); 2) Samples were loaded into Lysing Matrix E tubes and lysed with the FastPrep instrument (speed 5.5; 30-40 sec) (MP Biomedicals, Santa Ana, CA, USA), with lysis product centrifuged and loaded onto the NucleoSpin Filter; 3) DNase treatment was doubled in volume (190 $\mu$L) and time on-column (30 min).

#### Internal RNA standard
After RNA quantitation and before metatranscriptomic library prep, we also added a single stranded RNA internal standard using a linearized 917 bp *in vitro* transcribed fragment of the pTXB1 plasmid (New England Biolabs, Ipswich, MA, USA) [1, 2]. Though this internal standard was not used in the final analyses, reads belonging to the internal standard can be found in the final QC'ed reads. pTXB1 reads were removed from total RNA reads using bowtie2 (see Supplemental Table 6 for stats).

#### PacBio 16S rRNA gene survey
Extracted DNA was PCR-amplified (28-31 cycles) with standard bacterial 16S rRNA gene primers 27F/1492R [3]. The resulting amplicon product was then cleaned, ligated to a 17-mer barcoded adapter, amplified in a limited cycle PCR (20 cycles) to enrich the barcode-ligated product, and cleaned before sequencing at the University of Delaware DNA Sequencing and Genotyping Center (SMRTbell Template Prep Kit 1.0; PacBio RSII with P6-C4 chemistry and 6 hour movie) (PacBio, Menlo Park, CA, USA). Circular consensus reads with five full passes were used in the analysis (38,425 reads; mean length 1,322bp; mean number of passes 20.3x). Reads were assigned to 97% operational taxonomic units (OTUs) using QIIME [4]. One representative from each OTU was classified using SILVAngs [5] for classification. Zetaproteobacteria sequences were further classified into Zetaproteobacteria OTUs (ZOTUs) using ZetaHunter [6].

#### Metagenome and metatranscriptome sequencing
MG libraries were prepared using NEXTFLEX Rapid DNA-Seq Library Prep Kit for Illumina Platforms (Bioo Scientific Corp., Austin, TX, USA) with a target insert size of 250 bp, and were sequenced on an Illumina HiSeq 2500 (Illumina Inc., San Diego, CA, USA), with eight samples over two lanes of a paired-end 251-cycle run. Sample S9 was resequenced on a third of a lane on a paired-end 151-cycle run in an attempt to improve metagenome assembly quality. MT libraries were prepared from total RNA using the Ovation RNA-seq system V2 kit (NuGEN, Redwood City, CA, USA), which includes preferential priming of non-rRNA sequences, leading to ribosomal RNA sequence reduction. The prepped libraries were sequenced on the Illumina HiSeq 2500 platform, with 26 samples over one lane on a single-end 151-cycle run. An additional run on the HiSeq (1x51 bp; 5 samples; 1 lane) and the MiSeq (Nano; 2x151 bp; 3 samples) were

used in initial RNA recruitment, ribosomal reduction, and internal standard testing.

**Metagenome assembly, binning, and annotation**
Metagenomes were assembled using metaSPAdes with custom kmer settings: 21, 33, 55, 77, 99, 111, 127. In our assemblies, we were able to recover a better genome from S1_Zeta1 (79.4% of the overall community; up to 98% complete) by reducing its original coverage from 1,352x to 138x with a subassembly from 10% of the total reads (see discussion below in the section "Manual curation to locate cyc2 within the ZOTU2 S1_Zeta1 genome") (IMG Analysis Project ID Ga0261638). We also chose to re-assemble six previously published MG samples individually, as opposed to using the original coassembly to improve genome quality (IMG Analysis Project IDs Ga0256915, Ga0257019-Ga0257023) [7]. Other re-assemblies were also attempted, but only recovered bins with limited completeness (<31%) (data from [8, 9]).

**Zetaproteobacteria genome database and classification**
In addition to MAGs from this study and reassembled MAGs from Fullerton et al. [7], several published genomes were included in the Zetaproteobacteria genome database from SAGs [10, 11], MAGs [12–16], and isolates [17–24]. In total, 138 genomes were used either in the comparative genomic analysis, ribosomal protein phylogenetic tree, and/or had significant *cyc2* expression warranting discussion (see Supplemental Table 4).

To classify Zetaproteobacteria genomes, we first looked for an intact 16S rRNA gene within the genome, which could be classified to the ZOTU level using ZetaHunter [6]. All isolates and most SAGs possessed a 16S rRNA gene, while most MAGs did not. Following this, average amino acid identity (AAI) and average nucleotide identity (ANI) pairwise comparisons were calculated for all Zetaproteobacteria genomes to determine relatedness. AAI was calculated using the ruby gem "aai," written by Ryan Moore and designed for rapid calculation over multiple threads (github.com/mooreryan/aai). ANI was calculated using the OrthoANIu tool [25]. These data were combined with 16S ZOTU assignments to determine genome taxonomy. Finally, phylogenomic relationships of genomes were determined using a concatenated alignment of 12 ribosomal proteins. Phylogenetic trees from this alignment provide better resolution than 16S rRNA gene trees (see discussion below).

**Putative periplasmic cytochromes**
Putative periplasmic *c*-type cytochromes were identified in representative genomes from each ZOTU, and other representative Fe-oxidizing microbes, including *Sideroxydans lithotrophicus* ES-1, *Gallionella capsiferriformans* ES-2, *Ferriphaselus amnicola* OYT-1, *Acidithiobacillus ferrooxidans* ATCC 23270, and *Ferrovum myxofaciens* P3G. To identify these cytochromes *c*, genes were first filtered based on the CXXCH heme-binding motif, using a custom heme counter (github.com/seanmcallister/heme_counter). Next, a periplasmic signal sequence was identified in *c*-type cytochromes using SignalP 4.0 [26] and PSORTb [27]. In total, 52 different putative periplasmic cytochromes *c* (PCs) were identified. From these representatives, we used BLASTp to identify these genes in the rest of the Zetaproteobacteria genomes.

### Phylogenetic trees

The concatenated ribosomal tree was constructed using amino acid alignments from 12 ribosomal proteins (L5p, L6p, L14p, L15p, L17p, L18p, L24p, L30p, S4p, S5p, S11p, and S13p). These ribosomal proteins are found in most isolate and SAG reference genomes, allowing for the affiliation of phylogenetic groups within this tree with 16S rRNA gene-based ZOTUs. Genomes in the tree are reported in Supplemental Table 4. To construct the ribosomal protein tree, each protein was individually aligned using MUSCLE [28], manually masked to unambiguously-aligned positions, then concatenated into a single alignment file with all 12 ribosomal proteins. A maximum likelihood phylogenetic tree was built with RAxML version 7.2.8, using the GTR model with GAMMA approximation and 100 bootstraps [29].

Cyc2 phylogenetic trees were constructed by first identifying protein sequences with homology to Cyc2 in the NCBI and IMG databases using BLASTP (maximum e-value $1x10^{-5}$). This BLAST was repeated using queries representing all major Cyc2 clusters, including *Mariprofundus ferrooxydans* PV-1, *Gallionella capsiferriformans* ES-2, *Acidithiobacillus ferrooxidans* ATCC 23270, *Tenderia electrophaga*, and *Geobacter* sp. FRC-32. Identical sequences from these searches (from 2,413 results) were removed by clustering at 100% identity using CD-HIT, reducing the Cyc2 database to 977 protein sequences [30]. These sequences were aligned using MUSCLE [28]. Partial-length sequences were filtered from the database using this alignment, resulting in 897 sequences. The alignment was masked (>30% gap alignment columns removed) and a maximum likelihood phylogenetic tree was built using RAxML (392 positions, 100 bootstraps, CAT model of rate heterogeneity, JTT amino acid substitution model) [29]. Based on the resulting tree, highly sampled *Burkholderia* and *Xanthomonas* sequences were removed and 103 *cyc2* sequences recovered from MAGs from this study were added, resulting in a final phylogenetic tree with 634 full-length sequences. The resulting phylogenetic tree was colored and annotated using Iroki [31]. Separate trees of just the cytochrome region (61 amino acids) were also constructed. These show similar clustering patterns to the whole Cyc2 tree, but lack bootstrap support across the backbone, making it difficult to assess the tree topology.

### RNA read recruitment and expression estimates

We tested the fidelity of the read recruitment against MAG bins on their own and the entire MG assembly using ReadHog (github.com/Arkadiy-Garber/ReadHog) and found very little difference (median 1.1x higher recruitment in individual bin recruitment over whole assembly). This suggested that recruitment to the entire MG assembly was the best route for accurate expression estimates.

Only a subset of Zetaproteobacteria genomes could be normalized to constitutive gene expression (32% fewer Zetaproteobacteria genomes), due to expression of the constitutive genes being below detection. Given the low sequence depth of constitutively expressed genes in low abundance bins, constitutive normalized expression for some bins should be viewed with caution (see Supplemental Table 7). Bins with depth sufficient for a confident constitutive TPM are indicated where appropriate.

We were particularly interested in the accuracy of the read recruitment to the *cyc2* gene, and whether reads could accurately be assigned to different Zetaproteobacteria bins. Reads were recruited with a mean mismatch of 3.5 bp (maximum 28 bp). Though the cytochrome region of the *cyc2* gene has the highest conservation, there are more differences between genomes in the same ZOTU (37-68 differences, depending on ZOTU) than the maximum read mismatch. This suggests that expression estimates for the *cyc2* gene are accurate.

Raw gene counts were provided from the S6 metatranscriptome as a whole, and for each bin of interest to test for significant differential expression in DESeq [32]. However, no gene was significantly differentially expressed. This is likely due to the limited amount of replication in our environmental metatranscriptomes, which would result in low power for statistical testing.

### Supplemental Text

#### Microbial Fe mat descriptions from MG/MT samples

Fe mats sampled at Loihi, Mariana, and vents along the MAR had a variety of physical settings, textures, and dominant Fe biominerals morphologies. Mat textures at Loihi are the best described, ranging from cohesive curd mat material above direct venting orifices to veil mats draped over rocks near diffusive venting [33, 34]. Sample S1 was collected from the top 1 cm of surface mat material from a curd mat, directly above two venting orifices. Sample S6, also thought to be a curd mat, was collected using a suction sampler, removing the top layer of mat material poised on a rock above a venting fissure. Sample S19 was collected from a trough above an active vent using a scoop sampler, yielding deep mat material.

Fe mats at the Golden Horn chimney, Urashima vent field, Mariana Backarc, were less cohesive than the curd mats at Loihi, resembling a lighter texture similar to veil mats. The collected mats from Mariana all came from a single Fe mineralized chimney, reaching 7 m in height. Samples S7 (B4/B5), S8 (B1/B2), scoop8, and RNA later scoop 24 were all collected from the base and middle of the chimney, where cracks in the chimney structure led to diffuse venting where the Fe mats could grow.

Samples from the MAR were collected from three different venting regions, where tens of cm-scale tufts of Fe mats could be found at the periphery of hot black smoker vents. 664-BS3 and 664-scoop8 were collected from Rainbow, from the same Fe mound. 664-BS3 represents surface mat, while 664-scoop8 represents deeper mats and sediment. 664-BS3 was dominated by twisted stalks and “y-guys” as the Fe oxide biominerals. Sample 665-MMA12 was taken from the surface of an Fe mat dominated by sheaths and stalks at the TAG vents. Sample 667-BS4 represented a more veil-like surface mat sample, dominated by sheaths, though some stalks were also present.

**Manual curation to locate *cyc2* within the ZOTU2 S1_Zeta1 genome**
Many metagenome assembled genomes (MAGs) from ZOTU2 lack the *cyc2* gene, though its presence in 3 single cell genomes (SAGs) [10] and 6 low quality MAGs [7] suggest that the gene should be found within the lineage. Of the 9 high quality genomes considered for comparative genomics in this study, only 2 possessed *cyc2*: 1 from AB-133-M17 (SAG) and 1 from 481-BS4-Zeta6 (MAG from Fullerton et al. [7] reassembled for this study). The repeated absence of *cyc2* within ZOTU2 suggested that the genome region may be difficult to assemble automatically, potentially due to its high coverage. Other researchers have found that a coverage ~20-100X is more likely to produce more complete bins due to tradeoffs between coverage and sequence error/heterogeneity when organisms are oversampled [35, 36]. For this reason, we chose to focus manual curation on one near complete ZOTU2 genome in an attempt to recover *cyc2*.

Loihi metagenome sample 674-BM1-C3 was dominated by a single ZOTU2 bin (S1_Zeta1), accounting for 79% of the total binned average read coverage (50.3% including unbinned contigs). This bin was estimated by CheckM to be 92.7% complete (3.03% redundancy, 0% strain heterogeneity), yet lacked any protein BLAST hits to Cyc2. We used two methods to find the *cyc2* belonging to this bin: 1) we subsampled reads for a simplified assembly and 2) we used information from the assembly graph to locate *cyc2* connected with this dominant bin. Quadruplicate, randomly-sampled read subsets at 10%, 2%, and 1% of the total number of quality-controlled reads were independently assembled and binned to simplify the assembly of the dominant ZOTU2 bin (starting at 1,352x coverage). Subsampling in this way allowed us to increase the quality of S1_Zeta1, with a maximum completeness of 98.1% (2.27% redundancy, 0% strain heterogeneity; 10% subassembly), though *cyc2* was again not recovered. However, the subsampled assembly was helpful in recovering *cyc2* through use of the simplified assembly graph. Using BLASTp, 12 contigs were identified that had homology to Cyc2. All twelve of these contigs were found within one section of the assembly graph, connected through a minimal k-mer overlap of 21 bp to the assembly network containing the binned contigs of interest. No other bins were contained within this assembly graph network. Taking the section of the assembly graph containing these *cyc2* contigs, overlap consensus assembly produced five unique contigs. Four of these contigs had sufficient length to confirm their clustering within S1_Zeta1 using VizBin [37]. These four contigs were combined with the dominant subsampled bin and used in further analysis (S1_10_Zeta1 genome). Though requiring manual curation, these methods show that it is possible to recover *cyc2* in ZOTU2 genomes that originally lacked the gene after automated assembly.

**References**


1. Satinsky BM, Gifford SM, Crump BC, Moran MA. Use of internal standards for quantitative metatranscriptome and metagenome analysis. *Methods Enzymol* 2013; **531**: 237–250.
2. Cottrell MT, Kirchman DL. Transcriptional control in marine copiotrophic and oligotrophic bacteria with streamlined genomes. *Appl Environ Microbiol* 2016; **82**: 6010–6018.

3. Lane DJ. 16S/23S rRNA sequencing. *Nucleic Acid Tech Bact Syst* 1991; 115–175.
4. Caporaso JG, Kuczynski J, Stombaugh J, Bittinger K, Bushman FD, Costello EK, et al. QIIME allows analysis of high-throughput community sequencing data. *Nat Methods* 2010; **7**: 335–336.
5. Quast C, Pruesse E, Yilmaz P, Gerken J, Schweer T, Yarza P, et al. The SILVA ribosomal RNA gene database project: Improved data processing and web-based tools. *Nucleic Acids Res* 2013; **41**: D590–D596.
6. McAllister SM, Moore RM, Chan CS. ZetaHunter, a reproducible taxonomic classification tool for tracking the ecology of the Zetaproteobacteria and other poorly resolved taxa. *Microbiol Resour Announc* 2018; **7**: e00932-18.
7. Fullerton H, Hager KW, McAllister SM, Moyer CL. Hidden diversity revealed by genome-resolved metagenomics of iron-oxidizing microbial mats from Lōʼihi Seamount, Hawaiʼi. *ISME J* 2017; **11**: 1900–1914.
8. Singer E, Heidelberg JF, Dhillon A, Edwards KJ. Metagenomic insights into the dominant Fe(II) oxidizing Zetaproteobacteria from an iron mat at Lōʼihi, Hawaiʼi. *Front Microbiol* 2013; **4**: 52.
9. Meyer JL, Jaekel U, Tully BJ, Glazer BT, Wheat CG, Lin H-T, et al. A distinct and active bacterial community in cold oxygenated fluids circulating beneath the western flank of the Mid-Atlantic ridge. *Sci Rep* 2016; **6**: 22541.
10. Field EK, Sczyrba A, Lyman AE, Harris CC, Woyke T, Stepanauskas R, et al. Genomic insights into the uncultivated marine Zetaproteobacteria at Loihi Seamount. *ISME J* 2015; **9**: 857–870.
11. Scott JJ, Breier JA, Luther, III GW, Emerson D. Microbial iron mats at the Mid-Atlantic Ridge and evidence that Zetaproteobacteria may be restricted to iron-oxidizing marine systems. *PLoS One* 2015; **10**: e0119284.
12. Emerson JB, Thomas BC, Alvarez W, Banfield JF. Metagenomic analysis of a high carbon dioxide subsurface microbial community populated by chemolithoautotrophs and bacteria and archaea from candidate phyla. *Environ Microbiol* 2016; **18**: 1686–1703.
13. Probst AJ, Castelle CJ, Singh A, Brown CT, Anantharaman K, Sharon I, et al. Genomic resolution of a cold subsurface aquifer community provides metabolic insights for novel microbes adapted to high $CO_2$ concentrations. *Environ Microbiol* 2017; **19**: 459–474.
14. Probst AJ, Ladd B, Jarett JK, Geller-McGrath DE, Sieber CMK, Emerson JB, et al. Differential depth distribution of microbial function and putative symbionts through sediment-hosted aquifers in the deep terrestrial subsurface. *Nat Microbiol* 2018; **3**: 328–336.
15. Anderson RE, Reveillaud J, Reddington E, Delmont TO, Eren AM, McDermott JM, et al. Genomic variation in microbial populations inhabiting the marine subseafloor at deep-sea hydrothermal vents. *Nat Commun* 2017; **8**: 1114.
16. Lin W, Zhang W, Zhao X, Roberts AP, Paterson GA, Bazylinski DA, et al. Genomic expansion of magnetotactic bacteria reveals an early common origin of magnetotaxis with lineage-specific evolution. *ISME J* 2018; **12**: 1508–1519.
17. Emerson D, Rentz JA, Lilburn TG, Davis RE, Aldrich H, Chan CS, et al. A novel lineage of proteobacteria involved in formation of marine Fe-oxidizing microbial mat communities. *PLoS One* 2007; **2**: e667.

18. McAllister SM, Davis RE, McBeth JM, Tebo BM, Emerson D, Moyer CL. Biodiversity and emerging biogeography of the neutrophilic iron-oxidizing Zetaproteobacteria. *Appl Environ Microbiol* 2011; **77**: 5445–5457.
19. Fullerton H, Hager KW, Moyer CL. Draft genome sequence of Mariprofundus ferrooxydans strain JV-1, isolated from Loihi Seamount, Hawaii. *Genome Announc* 2015; **3**: e01118-15.
20. Mumford AC, Adaktylou IJ, Emerson D. Peeking under the iron curtain: development of a microcosm for imaging the colonization of steel surfaces by *Mariprofundus* sp. strain DIS-1, an oxygen-tolerant Fe-oxidizing bacterium. *Appl Environ Microbiol* 2016; **82**: 6799–6807.
21. Makita H, Tanaka E, Mitsunobu S, Miyazaki M, Nunoura T, Uematsu K, et al. *Mariprofundus micogutta* sp. nov., a novel iron-oxidizing zetaproteobacterium isolated from a deep-sea hydrothermal field at the Bayonnaise knoll of the Izu-Ogasawara arc, and a description of Mariprofundales ord. nov. and Zetaproteobacteria classis. *Arch Microbiol* 2017; **199**: 335–346.
22. Chiu BK, Kato S, McAllister SM, Field EK, Chan CS. Novel pelagic iron-oxidizing Zetaproteobacteria from the Chesapeake Bay oxic-anoxic transition zone. *Front Microbiol* 2017; **8**.
23. Mori JF, Scott JJ, Hager KW, Moyer CL, Küsel K, Emerson D. Physiological and ecological implications of an iron- or hydrogen-oxidizing member of the Zetaproteobacteria, *Ghiorsea bivora*, gen. nov., sp. nov. *ISME J* 2017; **11**: 2624–2636.
24. Beam JP, Scott JJ, McAllister SM, Chan CS, McManus J, Meysman FJR, et al. Biological rejuvenation of iron oxides in bioturbated marine sediments. *ISME J* 2018; **12**: 1389–1394.
25. Yoon S-H, Ha S, Lim J, Kwon S, Chun J. A large-scale evaluation of algorithms to calculate average nucleotide identity. *Antonie Van Leeuwenhoek* 2017; **110**: 1281–1286.
26. Petersen TN, Brunak S, von Heijne G, Nielsen H. SignalP 4.0: discriminating signal peptides from transmembrane regions. *Nat Methods* 2011; **8**: 785–786.
27. Yu NY, Wagner JR, Laird MR, Melli G, Rey S, Lo R, et al. PSORTb 3.0: Improved protein subcellular localization prediction with refined localization subcategories and predictive capabilities for all prokaryotes. *Bioinformatics* 2010; **26**: 1608–1615.
28. Edgar RC. MUSCLE: Multiple sequence alignment with high accuracy and high throughput. *Nucleic Acids Res* 2004; **32**: 1792–1797.
29. Stamatakis A. RAxML-VI-HPC: maximum likelihood-based phylogenetic analyses with thousands of taxa and mixed models. *Bioinformatics* 2006; **22**: 2688–2690.
30. Li W, Godzik A. Cd-hit: A fast program for clustering and comparing large sets of protein or nucleotide sequences. *Bioinformatics* 2006; **22**: 1658–1659.
31. Moore RM, Harrison AO, McAllister SM, Wommack KE. Iroki: automatic customization and visualization of phylogenetic trees. *bioRxiv* 2018; doi:10.1101/106138.
32. Anders S, Huber W. Differential expression analysis for sequence count data. *Genome Biol* 2010; **11**: R106.

33. Chan CS, McAllister SM, Leavitt AH, Glazer BT, Krepski ST, Emerson D. The architecture of iron microbial mats reflects the adaptation of chemolithotrophic iron oxidation in freshwater and marine environments. *Front Microbiol* 2016; **7**: 796.
34. McAllister SM, Moore RM, Gartman A, Luther GW, Emerson D, Chan CS. The Fe(II)-oxidizing Zetaproteobacteria: Historical, ecological, and genomic perspectives. *FEMS Microbiol Ecol* 2019; fiz015.
35. Brown CT, Howe A, Zhang Q, Pyrkosz AB, Brom TH. A reference-free algorithm for computational normalization of shotgun sequencing data. *arXiv* 2012; arXiv:1203.4802v2.
36. Coutinho FH, Silveira CB, Gregoracci GB, Thompson CC, Edwards RA, Brussaard CPD, et al. Marine viruses discovered via metagenomics shed light on viral strategies throughout the oceans. *Nat Commun* 2017; **8**: 15955.
37. Laczny CC, Sternal T, Plugaru V, Gawron P, Atashpendar A, Margossian HH, et al. VizBin - an application for reference-independent visualization and human-augmented binning of metagenomic data. *Microbiome* 2015; **3**: doi:10.1186/s40168-014-0066-1.

A) Loihi Seamount (2013)

Sample S1
J2-674-BM1-C3
Pohaku (Mkr 57)
Syringe
RNALater

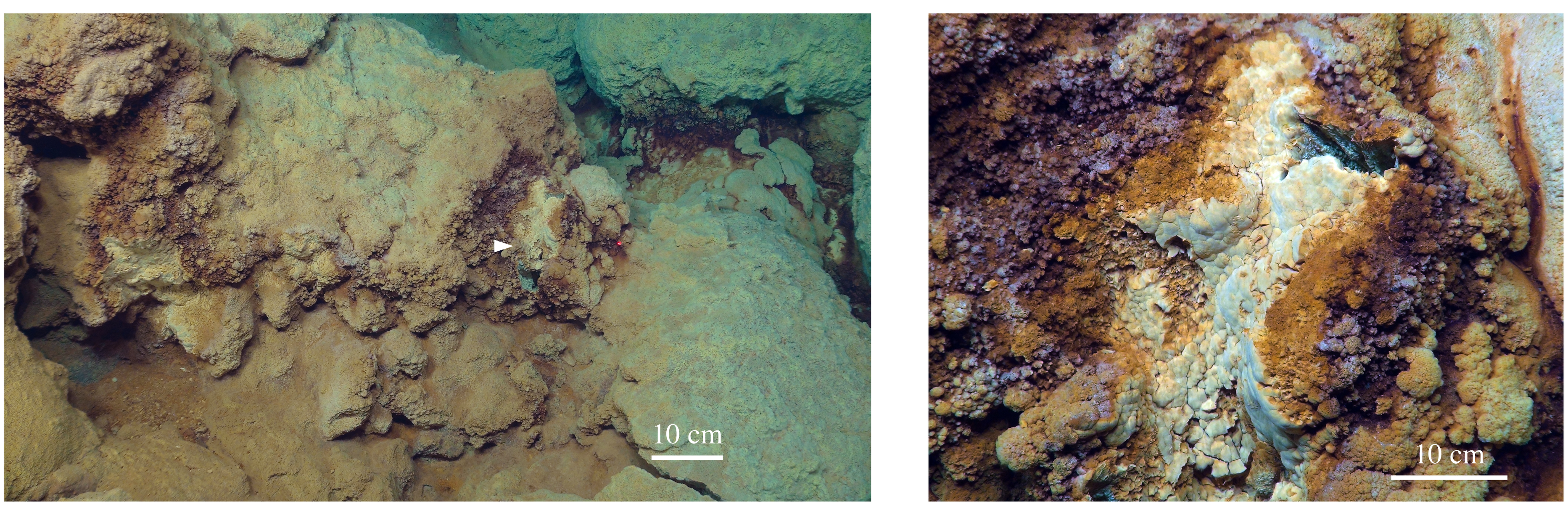


Sample S6
J2-677-SSyellow
Spillway (Mkr 34)
Suction sampler
Onboard experiment

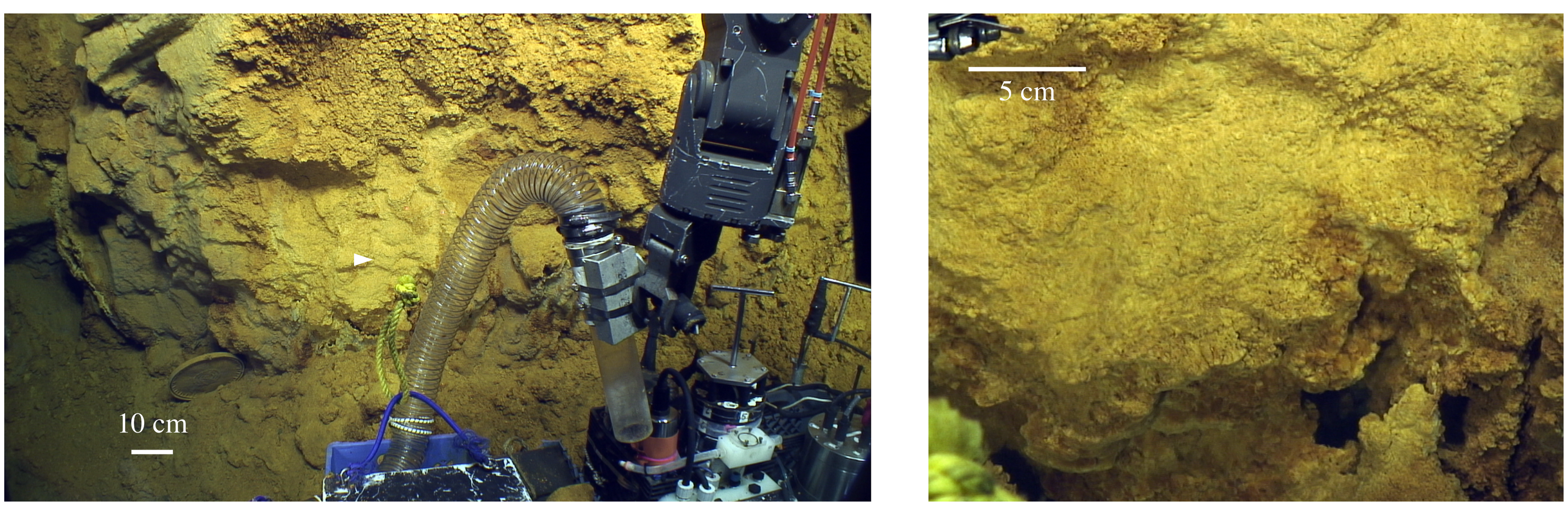


Sample S19
J2-675-SC9
Crop Circle (U Mkr 31)
Scoop
RNALater

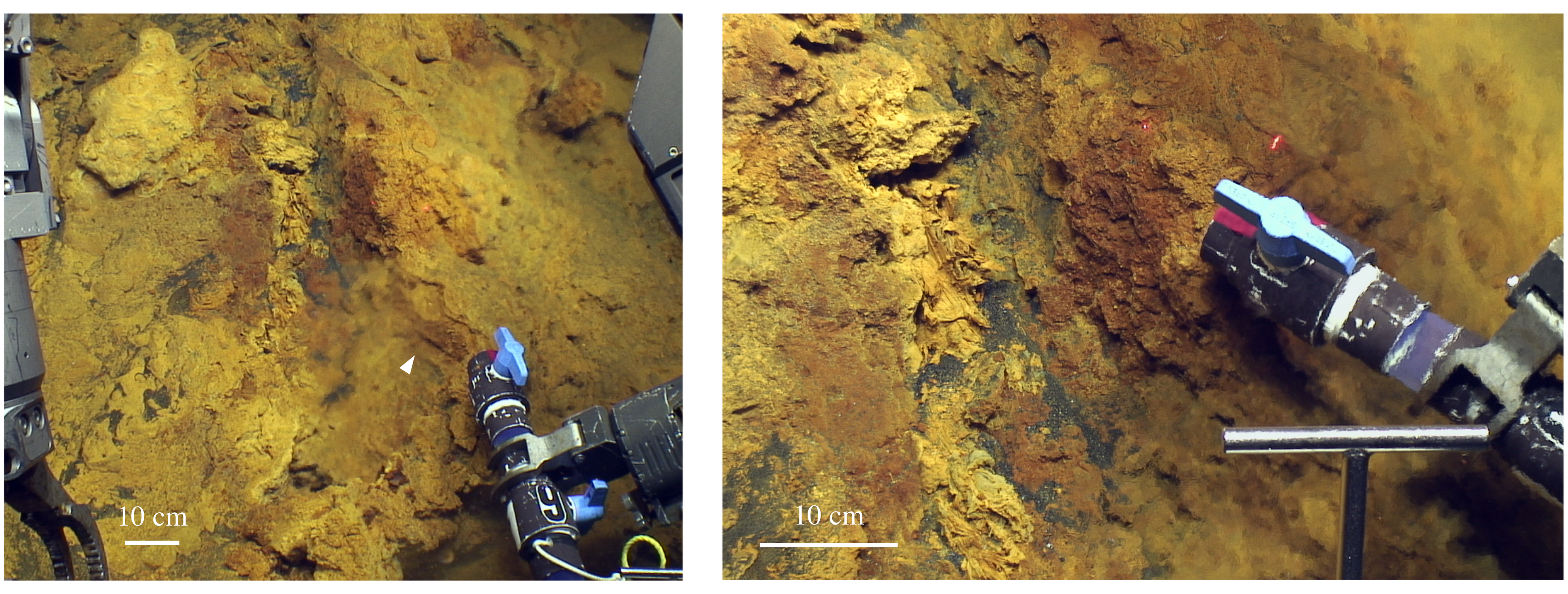


B) Mid-Atlantic Ridge (2012)

J2-664-BS3/
J2-664-SC8
Rainbow vent field
Syringe/Scoop
Untreated

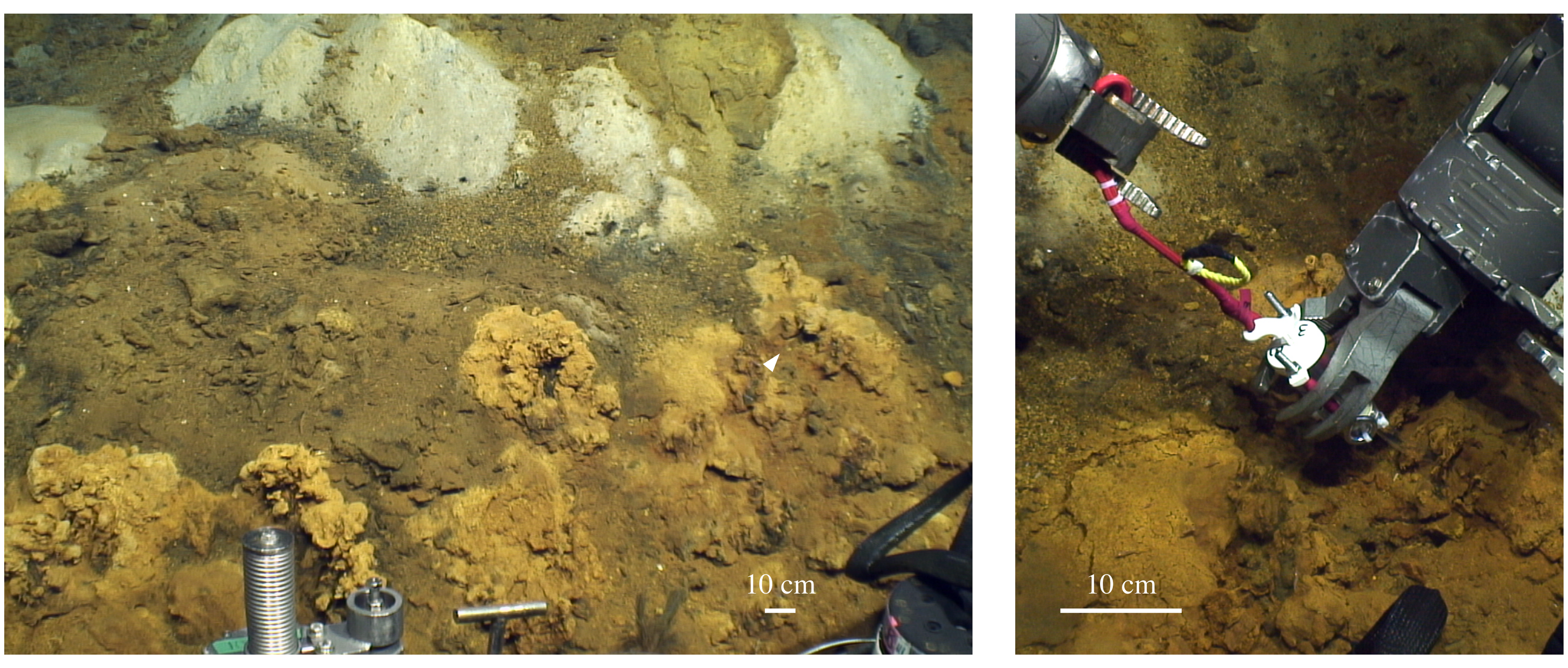


J2-665-MMA12
TAG vent field
Syringe
RNALater

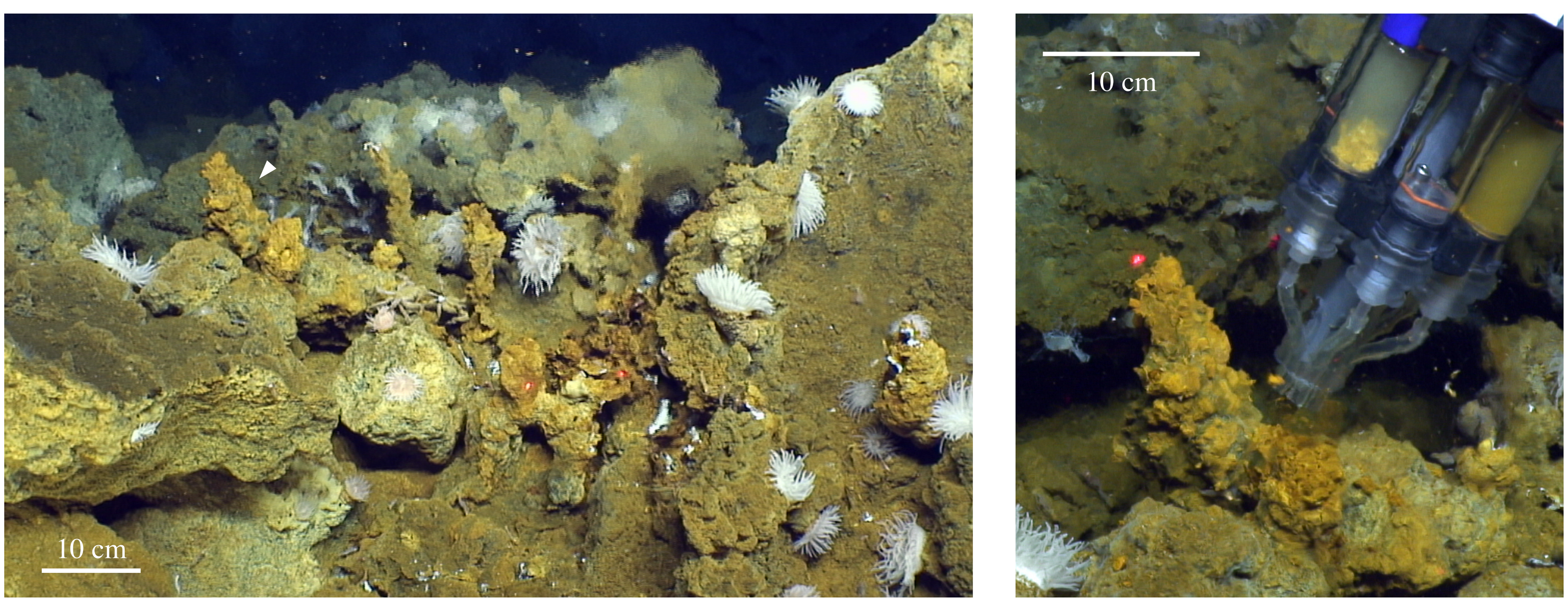


J2-667-BS4
Snake Pit vent field
Syringe
RNALater

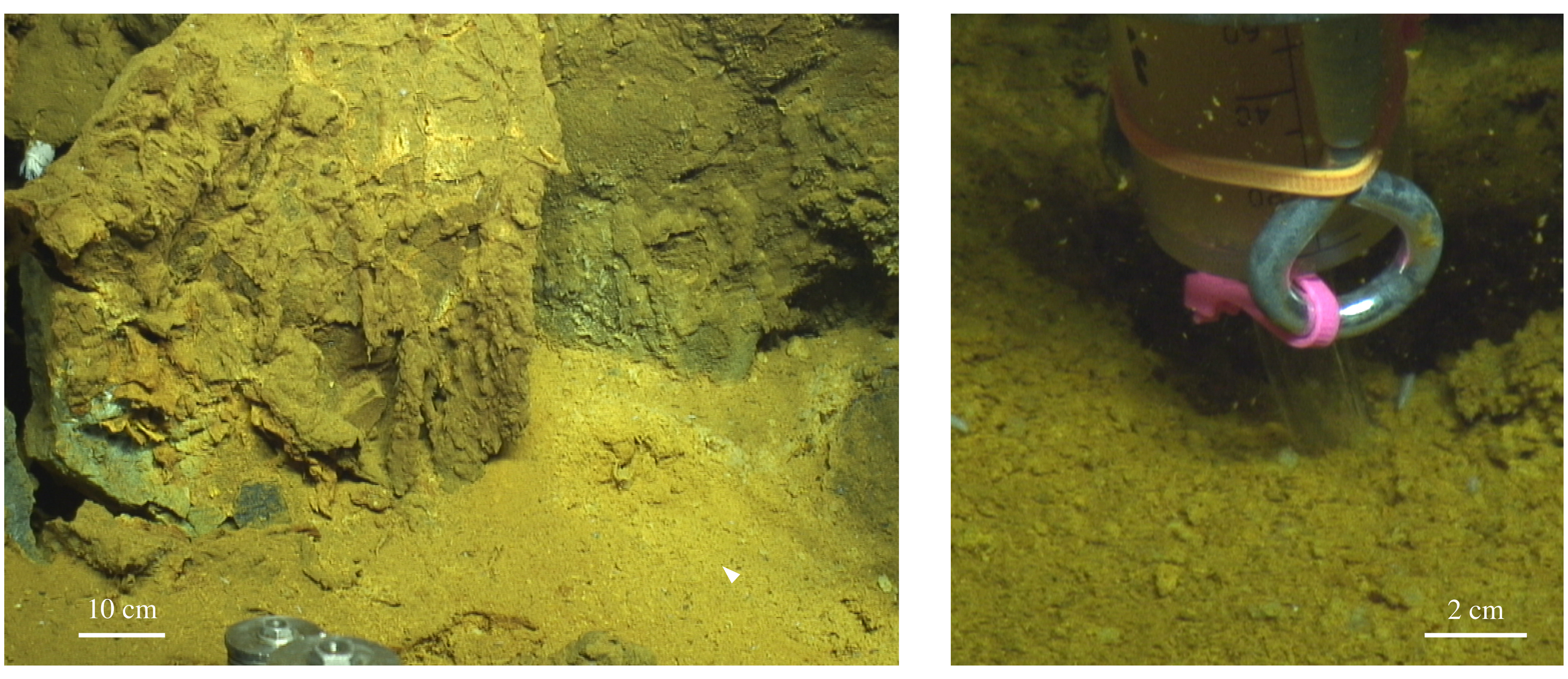


C) Mariana Backarc (2014)

Sample S7-B4/B5
J2-801-BM1-B4/B5
Urashima vent field
Golden Horn Chimney
Syringe
RNALater

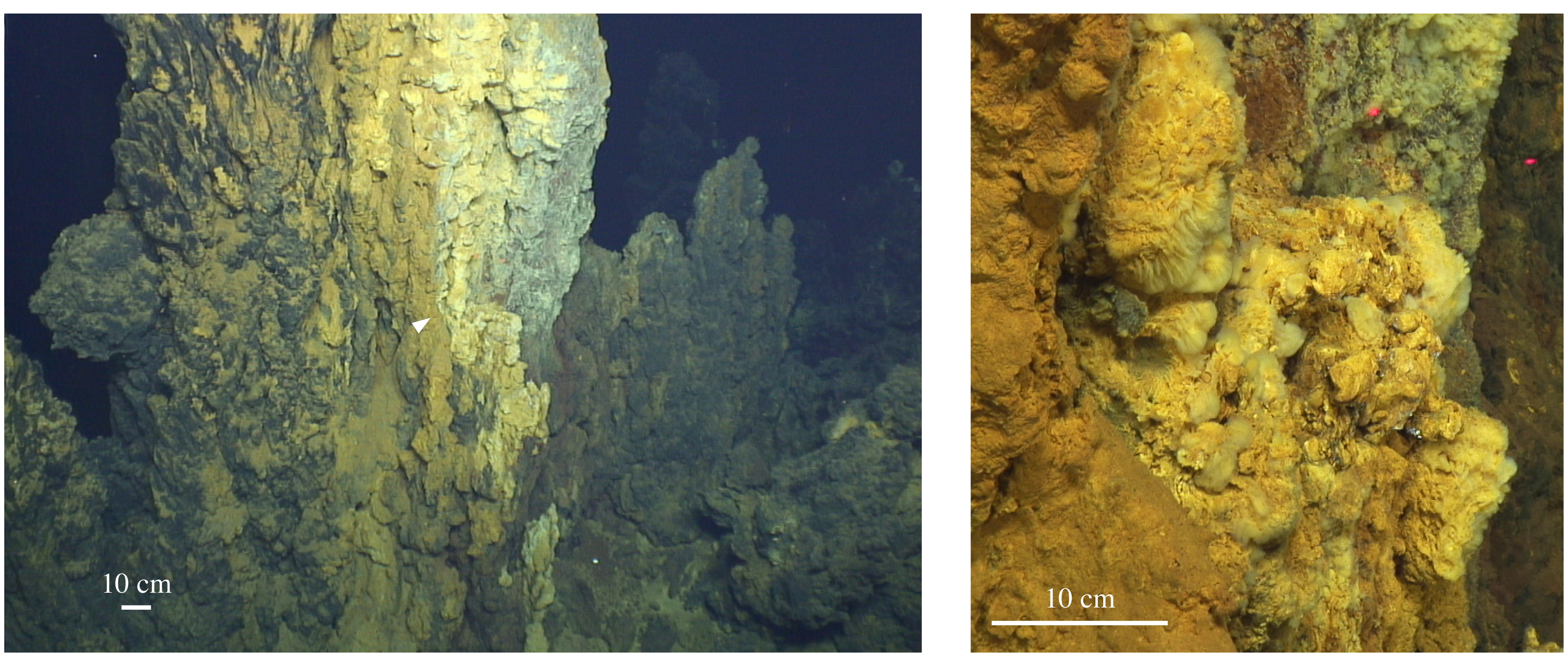


Sample S8-B2/B3
J2-801-BM1-B2/B3
Urashima vent field
Golden Horn Chimney
Syringe
RNALater

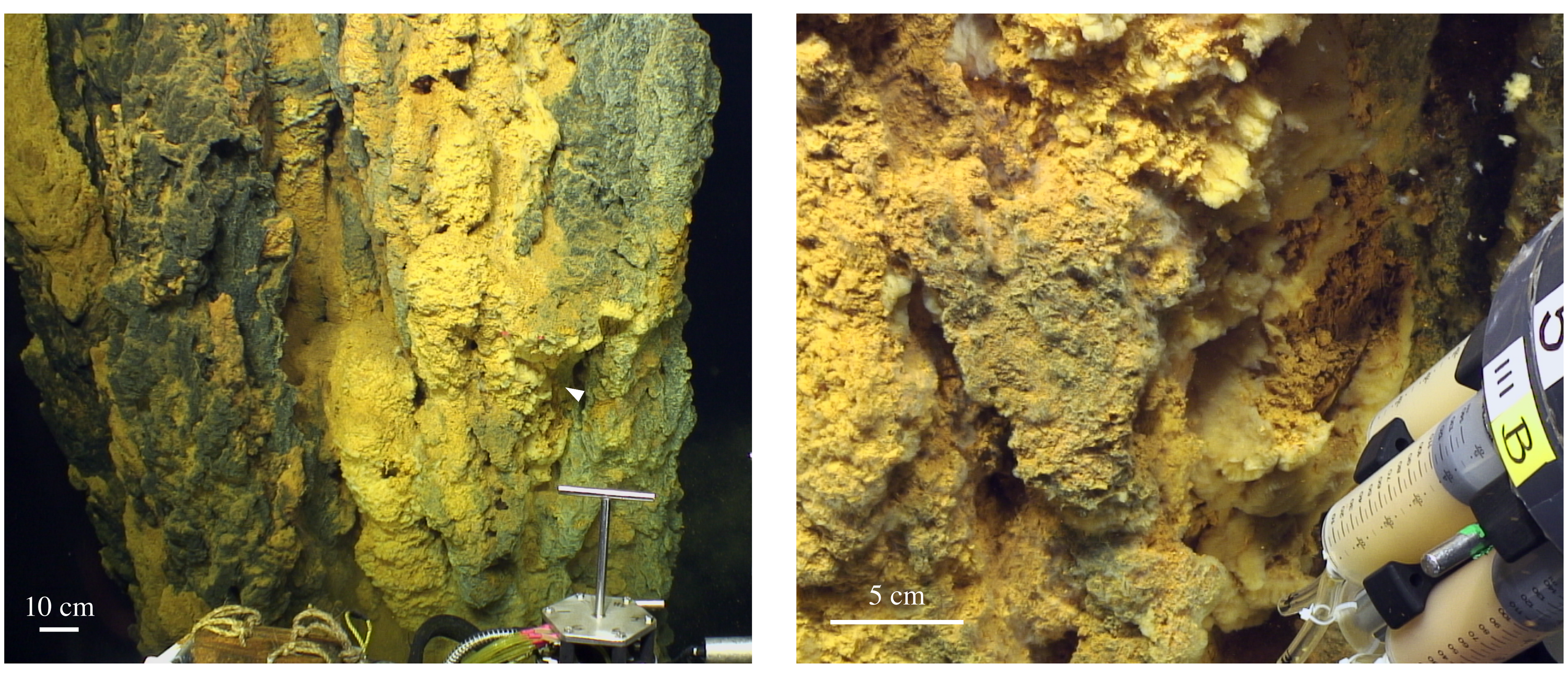


Sample S9
J2-801-SC8
Urashima vent field
Golden Horn Chimney
Scoop
Onboard experiment

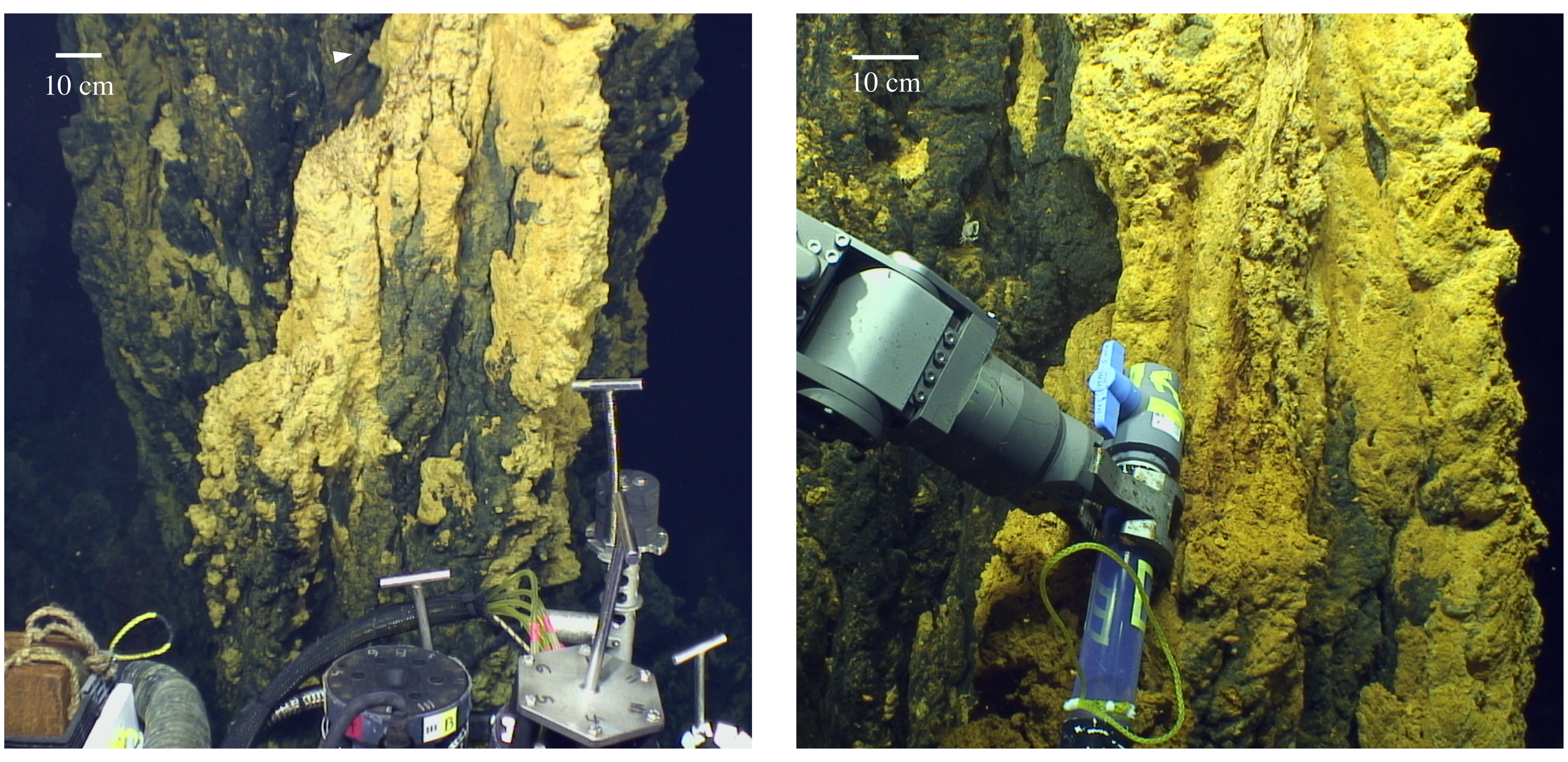


**Supplemental Figure 1.** Photographs of sampling locations for samples chosen for metagenome and metatranscriptome sequencing from (A) Loihi Seamount, (B) Mid-Atlantic Ridge, and (C) Mariana Backarc. Images on the left show context for the specific sampling location, which is marked by an arrowhead and depicted on the right.

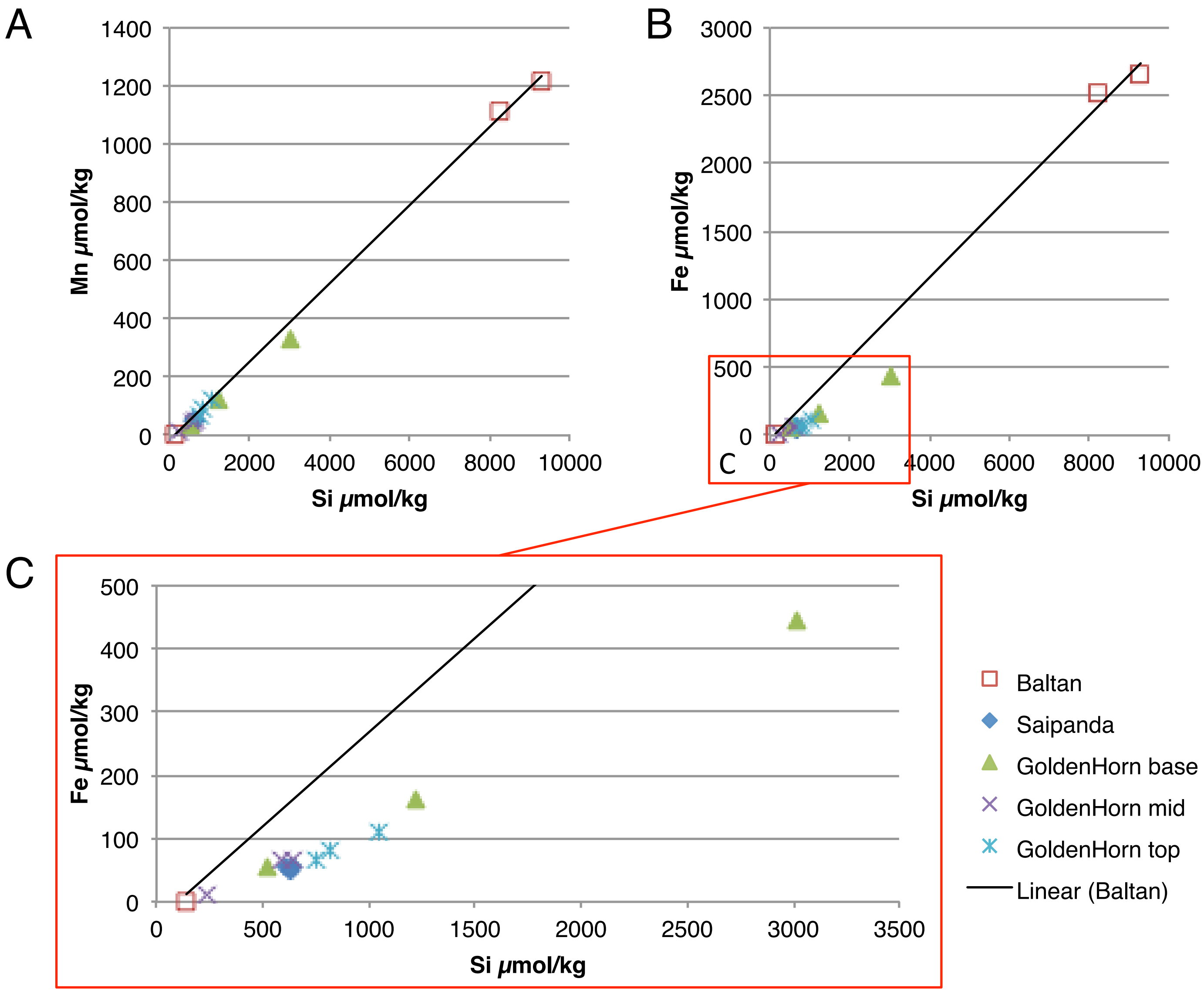


**Supplemental Figure 2.** Urashima Vent Field (Mariana) geochemistry, plotted to highlight mixing of a single endmember with seawater. Mn and Si are conservative, or unreactive, during mixing (A). Total dissolved Fe is depleted in low-temperature fluids compared to conservative mixing (B, enlarged in C). This is particularly pronounced within the Golden Horn Chimney Fe mats. Baltan is a high-temperature vent in the Urashima Vent Field. Saipanda is another low temperature vent that was not sampled for this study.

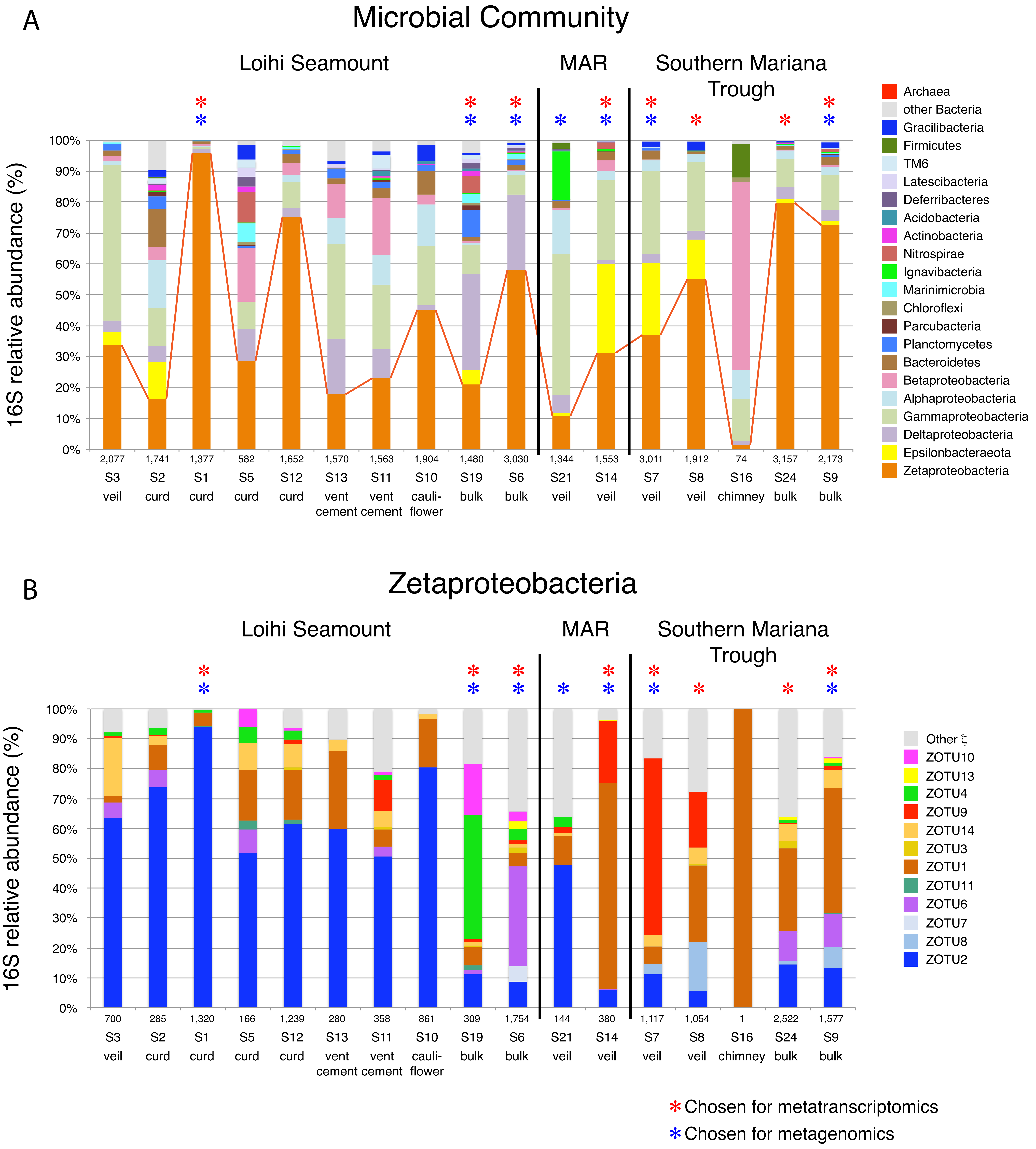


**Supplemental Figure 3.** PacBio 16S rRNA gene survey of Bacteria (A) and Zetaproteobacteria (B) microbial communities from Fe mats at Loihi Seamount, the Mid-Atlantic Ridge, and Southern Mariana Trough. The abundance of Zetaproteobacteria are highlighted (A). Blue asterisks denote samples chosen for metagenomics. Red asterisks denote samples chosen for metatranscriptomics. Numbers at the bottom of the bar charts denote the number of total 16S rRNA gene sequences sampled. Sample short name and Fe mat type are also given.

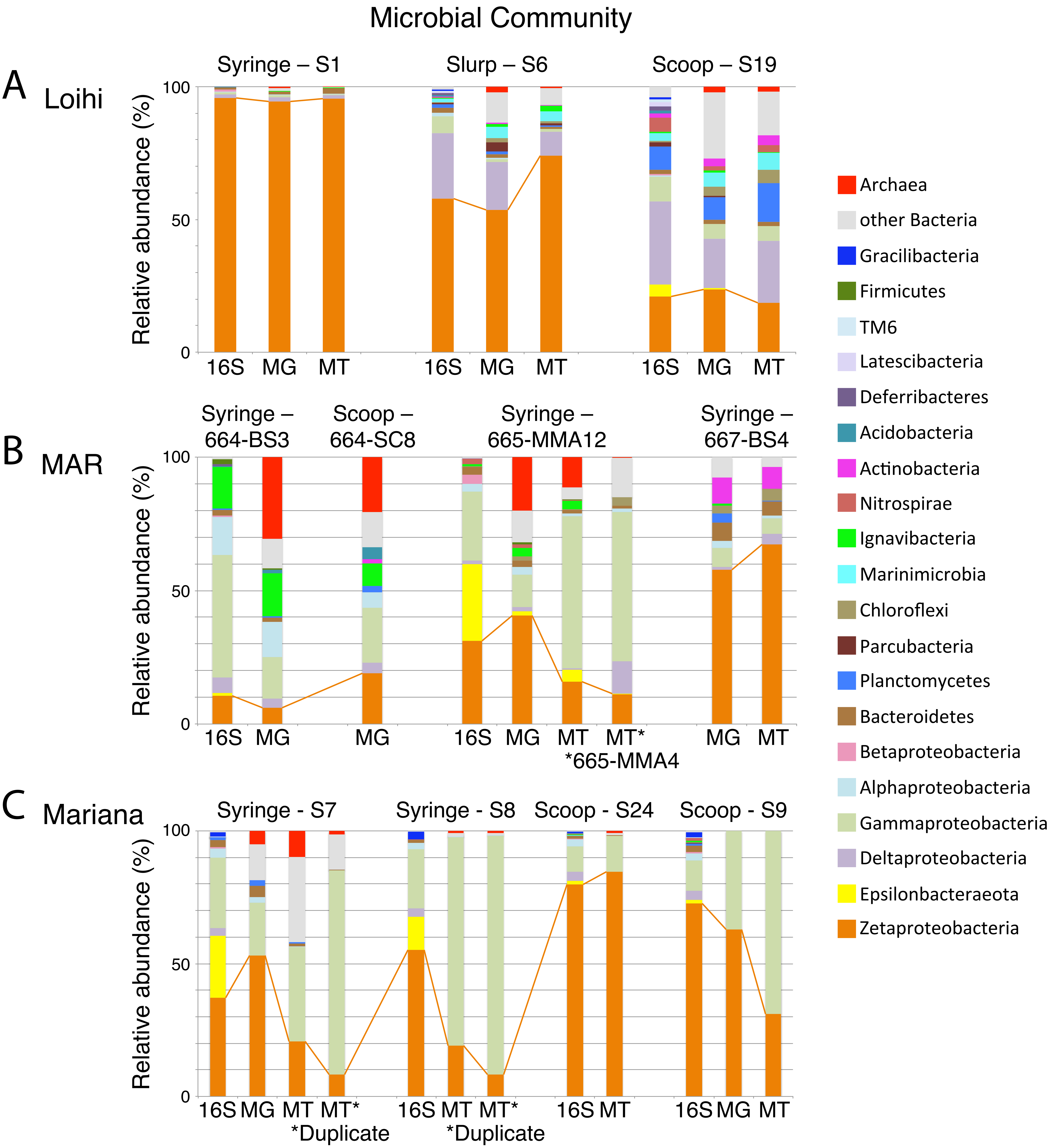


**Supplemental Figure 4.** Comparison of 16S rRNA gene, metagenome, and metatranscriptome relative abundance for the microbial communities at Loihi Seamount (A), the Mid-Atlantic Ridge (B), and Southern Mariana Trough (C). 16S rRNA gene plots represent the bacterial population only. The relative abundance of the Zetaproteobacteria are tracked for samples from the same Fe mat location and/or for MT samples mapped to the same metagenomes. Asterisks show MT samples that were mapped to a reference MG from a different sample.

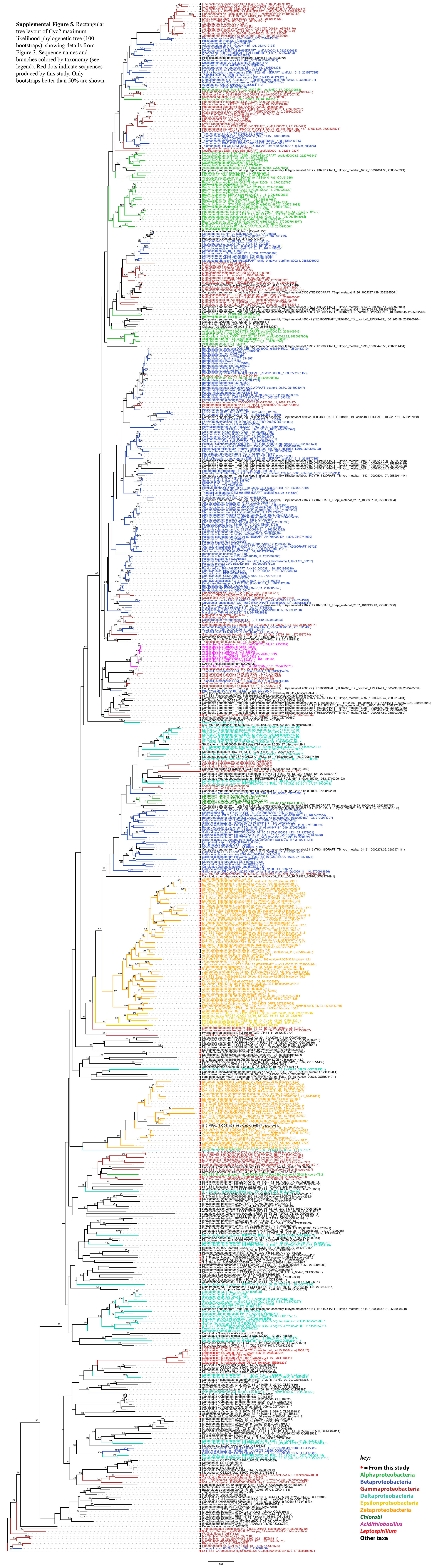


**Supplemental Figure 5.** Rectangular tree layout of Cyc2 maximum likelihood phylogenetic tree (100 bootstraps), showing details from Figure 3. Sequence names and branches colored by taxonomy (see legend). Red dots indicate sequences produced by this study. Only bootstraps better than 50% are shown.

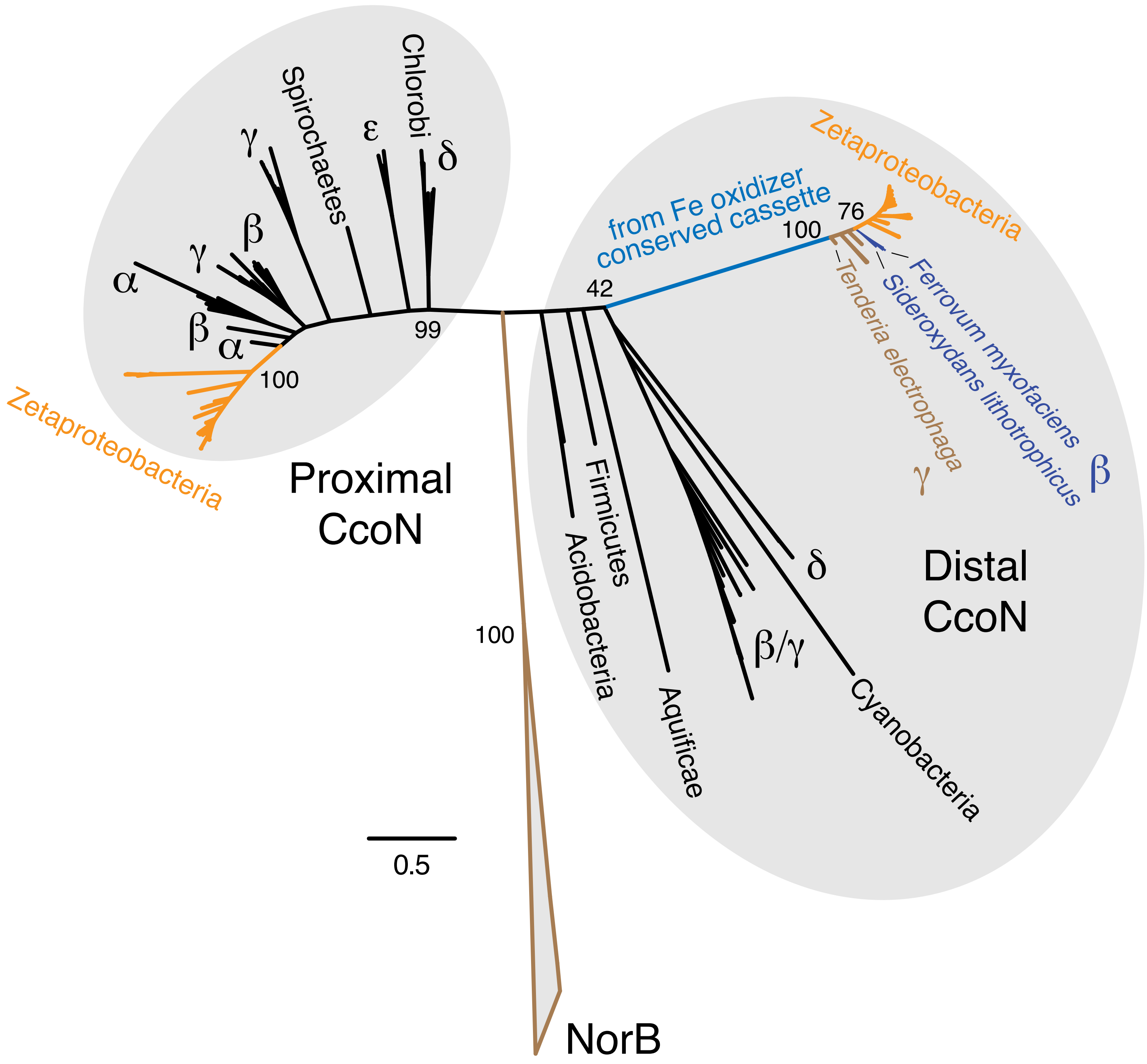


**Supplemental Figure 6.** Maximum likelihood phylogenetic tree of CcoN, showing that the Zetaproteobacteria possess two forms of *cbb*$_3$-type cytochrome c oxidase: proximal and distal. Only the distal version is found within the conserved cassette discovered by Field *et al.* (2015).

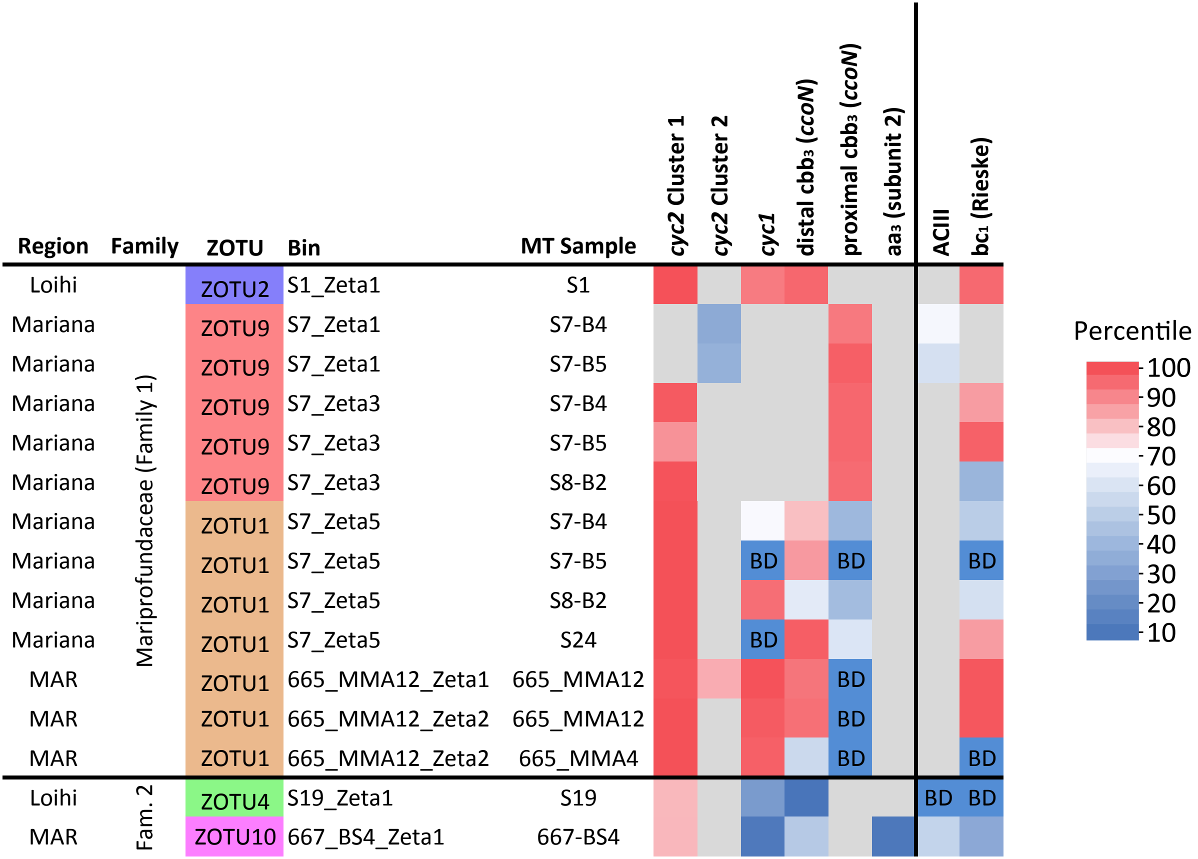


**Supplemental Figure 7.** Heatmap showing the percentile expression for key genes in the Fe oxidation pathway including genes thought to be involved in electron transport from Fe(II) to $O_2$ and from Fe(II) to the quinone pool for reverse electron transport (RET).

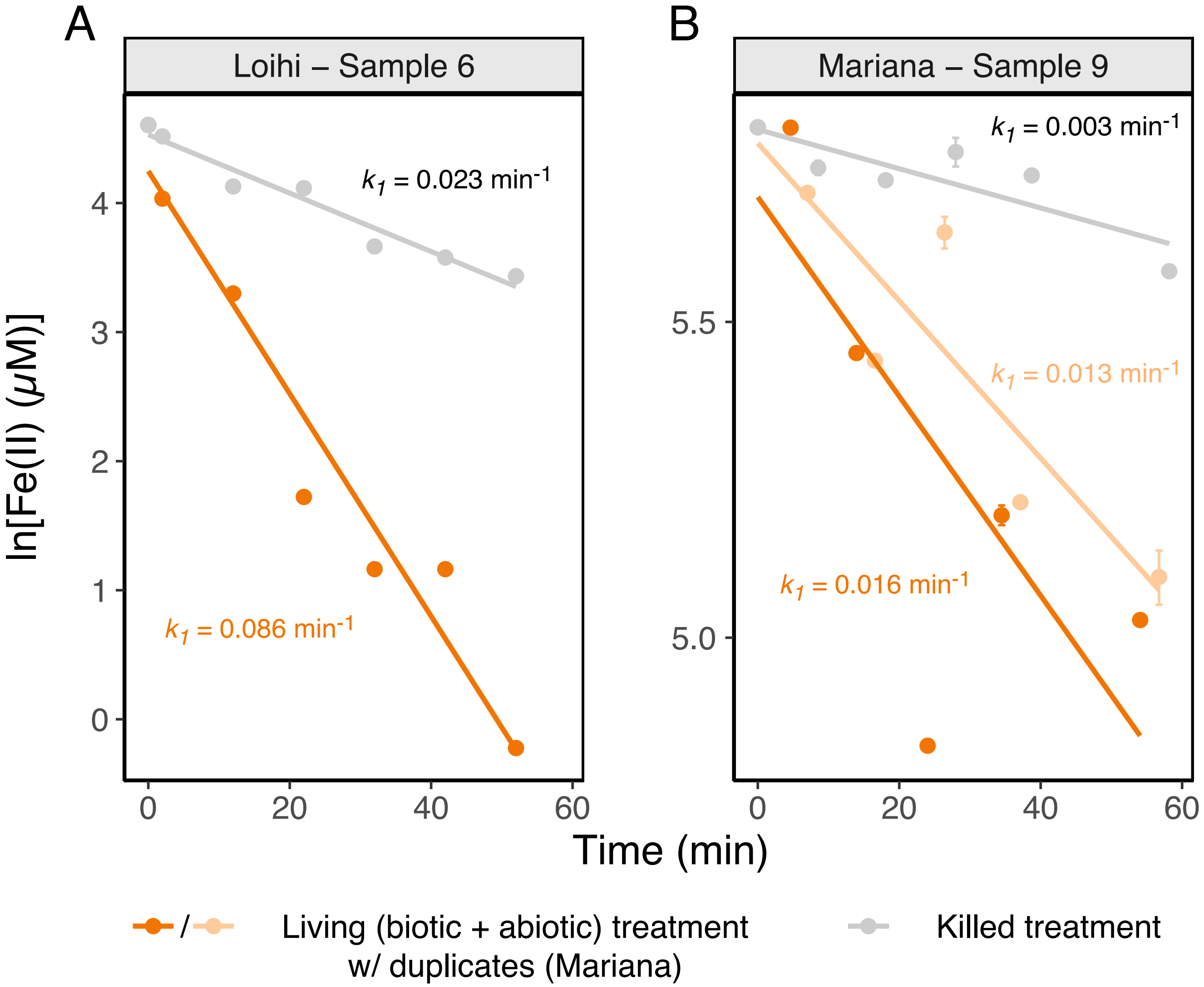


**Supplemental Figure 8.** Plot of Fe(II) addition experiment results from Loihi and Mariana Fe mats, showing a faster living (orange; total) versus killed (black; abiotic-only) Fe oxidation rate. Fe(II) was added to dormant Fe mat samples at zero min. Pseudo-first-order rate constants were calculated from the log-linear best fit from each experimental condition.

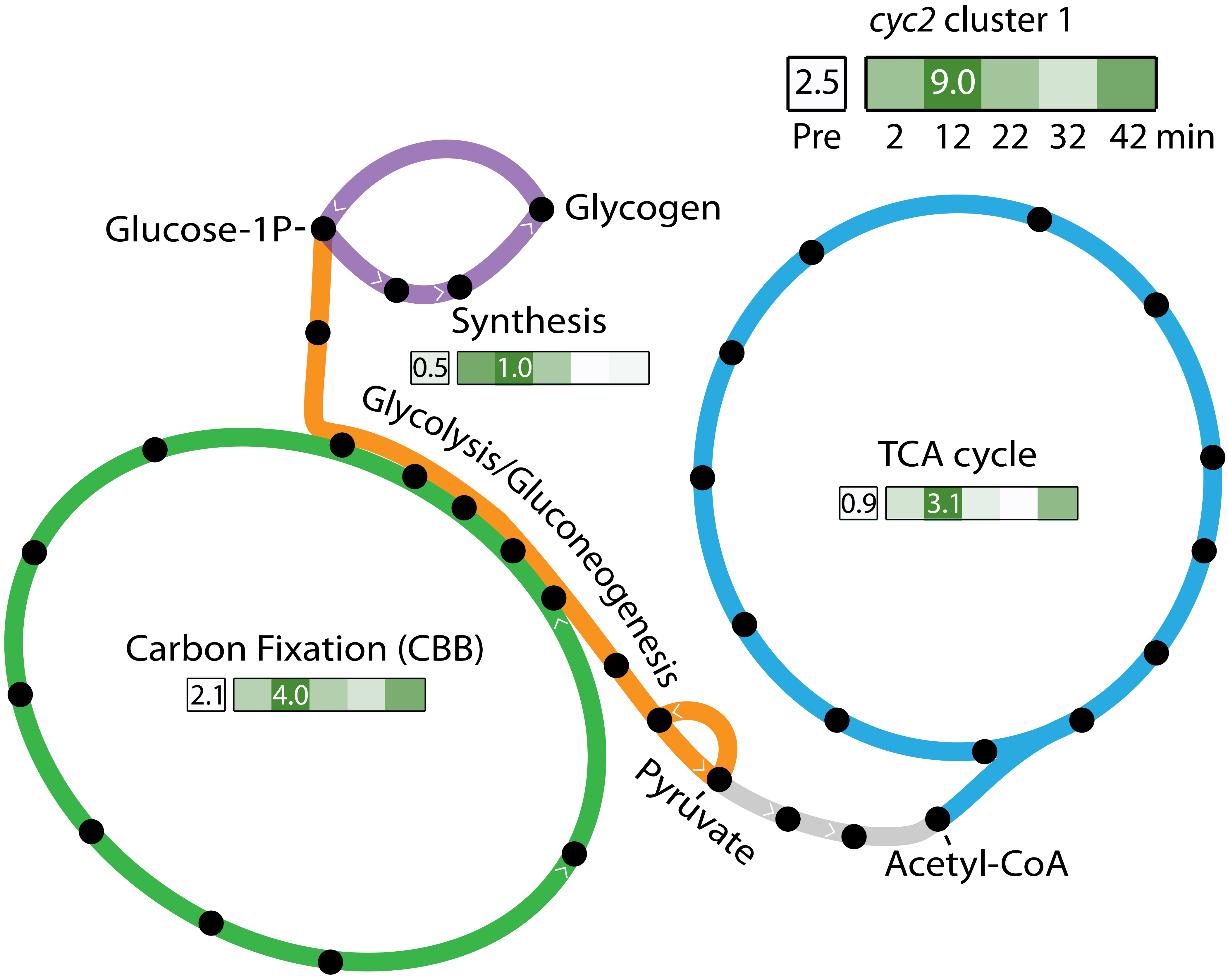


**Supplemental Figure 9.** Partial model of carbon metabolism in the S6_Zeta1 ZOTU6 genome. Changes in constitutive normalized expression are visualized as a max-normalized average of unique genes in each component of the model, with expression TPM pre-Fe(II) addition and at the highest point after Fe(II) addition indicated. Expression changes for the putative Fe oxidase, *cyc2*, is shown for comparison.
